# CRISPR–Cas9 Editing of Myeloid Lectins Reveals Target-Dependent Transcriptional Remodeling in Tumor-Associated Macrophages

**DOI:** 10.64898/2026.10.02.756241

**Authors:** Gonçalo Trindade, Miguel Pinto, Giacomo Domenici, Jose Escandell, Catarina Brito

**Affiliations:** iBET, Instituto de Biologia Experimental e Tecnológica, Oeiras, Portugal; Instituto de Tecnologia Química e Biológica António Xavier, Universidade NOVA de Lisboa, Oeiras, Portugal

## Abstract

Tumor-associated macrophages (TAMs) integrate multiple microenvironmental signals, including lectin-mediated recognition of tumor-associated glycans. Although several myeloid lectins have been implicated in tumor–macrophage interactions, their contributions to immunosuppressive TAM phenotypes remain poorly defined. We implemented a CRISPR–Cas9 ribonucleoprotein (RNP) workflow in primary human monocytes to enable comparative loss-of-function interrogation of five potentially immunosuppressive lectins we previously reported. CRISPR–Cas9 RNP delivery conditions were benchmarked in freshly isolated monocytes using *CD14* as a technical target, followed by sgRNA editing efficiency screening for *CLEC4E*, *CLECcA*, *CD20S*, *SELL*, and *SIGLEC9* in HeLa S3 cells. Selected guides were applied to primary monocytes before co-culture in a 3D model with BT474 breast cancer cells. Editing was quantified by Sanger-sequencing deconvolution, while co-culture-derived CD45⁺ macrophages were isolated by fluorescence-activated cell sorting and characterized by bulk RNA sequencing. All five lectins were successfully perturbed, although gene editing efficiency remained target- and donor-dependent. Transcriptomic analysis revealed target-specific reprogramming and recurrent immunometabolic responses. mTORC1 signaling was consistently enriched, together with the remodeling of cholesterol homeostasis, fatty acid metabolism, and oxidative phosphorylation. *CLEC6A* deletion achieved attenuation of hypoxia-, angiogenesis-, and TNF/NF-κB-associated programs. *CD20S* and *SIGLECS* deletion similarly reduced hypoxia- and angiogenesis-associated signatures while promoting lipid and oxidative metabolism. *CLEC4E* perturbation induced broad remodeling of lipid and oxidative metabolism and stress-response, despite comparatively limited editing efficiency. Comparative lectin perturbation thus identified *CLEC6A* as the strongest candidate for further mechanistic investigation, while *CLEC4E* showed broad transcriptional remodeling despite lower editing efficiency, supporting its prioritization for improved genetic interrogation. Responses to *CD20S* and *SIGLECS* targeting further support tumor glycan–myeloid signaling as a regulator of TAM state. Overall, lectin perturbation redirected TAM transcriptional states through distinct immunometabolic programs rather than a uniform pro-inflammatory polarization, highlighting lectin-dependent immunometabolic regulation as a promising axis for further investigation.

## Introduction

Tumor-associated macrophages (TAMs) are highly plastic components of the tumor microenvironment, where tumor-derived signals can shape TAM states associated with immune regulation, angiogenesis, tissue remodeling, and tumor progression.^1^ Consistent with this plasticity, studies of human tumors have shown that monocytes and macrophages undergo extensive transcriptional reprogramming and acquire heterogenous TAM phenotypes, emphasizing that their functional states extend beyond a simple M1/M2 classification.^1–3^ This phenotypic diversity reflects the integration of multiple environmental cues, including soluble factors such as cytokines and cell-surface recognition systems.^4^ Among these, tumor-associated glycosylation has emerged as an important layer of immune regulation.^5^ Tumor cells frequently display altered glycosylation of cell-surface and secreted molecules, including changes in sialylation and fucosylation, which can modify recognition by glycan-binding lectin receptors expressed on immune cells.^6,7^

Emerging evidence implicates several myeloid lectin receptors in tumor–macrophage communication.^5,7^ In breast cancer, fucosylated tumor-associated ligands can engage DC-SIGN (*CD209*) on TAMs and promote proangiogenic responses, while hypersialylated MUC1 can engage Siglec-9 (*SIGLEC9*) and influence monocyte differentiation toward TAM states.^8,9^ Mincle (*CLEC4E*) has similarly been implicated in macrophage-dependent tumor immune suppression and tumor-associated lipid sensing, whereas Dectin-2 (*CLEC6A*) has been shown to participate in macrophage recognition and phagocytosis of tumor cells.^10–12^ L-selectin (*SELL*), although less directly studied in TAM-state regulation, is a major regulator of leukocyte adhesion and trafficking and may therefore influence monocyte recruitment to tumor tissues.^13,14^ Collectively, these observations position lectins as an emerging regulatory layer in TAM biology, although the receptor-specific contribution of individual lectins to tumor-conditioned macrophage states remains unresolved.^15^

Addressing this gap requires moving beyond receptor expression, which alone does not establish a functional contribution to the TAM state. Genetic loss-of-function therefore provides a direct approach to interrogate receptor function.^16^ Primary human monocytes, however, remain technically challenging targets for genetic manipulation, as perturbation strategies must achieve sufficient efficiency while preserving cell viability, differentiation capacity, and downstream functional responsiveness.^17,18^ CRISPR–Cas9 ribonucleoprotein (RNP) delivery using nucleofection provides a non-viral approach in which pre-assembled Cas9–sgRNA complexes are introduced directly into cells, avoiding the need for continuous Cas9 expression from plasmid or viral vectors and resulting in transient exposure to the editing machinery and reduced off-target mutations while generating stable genomic modifications.^17,19^ In the absence of a repair template, Cas9-induced DNA double-strand breaks are predominantly resolved through non-homologous end joining (NHEJ) and related end-joining pathways, generating insertions and deletions (indels), including frameshift mutations that can disrupt protein-coding sequences.^16^ Previous studies have demonstrated the feasibility of RNP-based editing in primary human monocytes while preserving macrophage differentiation and functional responsiveness; however, relatively few workflows have been described, with studies employing distinct nucleofection conditions and no broadly standardized strategy.^17,18^ Nevertheless, the demonstrated feasibility of RNP-based editing provides an opportunity to combine receptor-level genetic perturbation with tumor-conditioning models and transcriptomic profiling, thereby distinguishing receptor-specific effects from biological responses shared across different myeloid sensing pathways.

Building on prior candidate-prioritization analyses of lectin expression during tumor-conditioned macrophage differentiation, and emerging evidence implicating lectin-mediated pathways in tumor-associated myeloid regulation, we selected *CLEC4E*, *CLECcA*, *CD20S*, *SELL*, and *SIGLEC9* for functional interrogation.^20^ Thus, we implemented a CRISPR-Cas9 RNP workflow for comparative genetic perturbation in freshly isolated primary human monocytes, incorporating nucleofection benchmarking, sgRNA screening, and sequence-based assessment of editing efficiency. The selected lectins were subsequently perturbed during macrophage differentiation in a BT474 breast cancer– monocyte co-culture model, followed by recovery of CD45⁺ macrophages for transcriptomic analysis. This framework enabled comparative assessment of the transcriptional effects of individual lectin perturbations, examining both target-specific responses and convergence across broader biological programs.

## Materials and Methods

### Cell Line Culture

BT474 and HeLa S3 cell lines were acquired from ATCC and cultured in RPMI 1640 or DMEM medium (Gibco), respectively, both supplemented with 10% (v/v) FBS (Gibco). Cells were sub-cultured twice a week at a seeding density of 30,000 cell/cm^2^. After thawing, cell lines were sub-cultured at least twice before being used for downstream assays. Cell cultures were maintained in a humidified incubator at 37 °C and 5% CO_2_.

### Human Peripheral Blood-Derived Monocyte Isolation

Human buffy coats from healthy donors were obtained from the Portuguese Blood Bank (Instituto Português do Sangue e da Transplantação – IPST). Isolation of PBMCs from buffy coats and then monocytes from PBMCs was performed as reported previously.^21,22^ Briefly, buffy coats were diluted 1:1 with Dulbecco’s Phosphate-Buffered Saline (DPBS) with 2% (v/v) FBS and 2 mM EDTA (Promega). Diluted buffy coats were processed by density centrifugation with Lymphoprep™ on SepMate™-50 (StemCell Technologies), and the peripheral blood-derived mononuclear cell (PBMCs) fraction was recovered. Subsequent washing steps and incubation with ACK Lysis Buffer (Gibco) were performed on PBMCs to minimize contamination with platelets and red blood cells. Monocytes (CD14^+^CD16^-^) were isolated through immunomagnetic negative selection, using the EasySep Human Monocyte Isolation Kit (StemCell Technologies). Monocytes were used directly either for macrophage differentiation in 2D cultures or co-culture with cancer cells in 3D spheroids.

### Primary Monocyte Culture

Monocyte monocultures were maintained in basal medium composed of RPMI 1640 medium without Phenol Red (Gibco), with 10% (v/v) FBS, 1% (v/v) Penicillin-Streptomycin (P/S), and supplemented with 50 ng/mL Macrophage Colony-Stimulating Factor (M-CSF) (Peprotech, #300-25). Monocytes were plated at 300,000 cell/cm^2^ and cultured up to 7 days in a humidified incubator at 37 °C and 5% CO_2_. For differentiation of M0, M1 or M2 states, at day 5 medium was replaced with basal medium with no supplementation for M0, supplemented with 10 ng/mL lipopolysaccharide (LPS, Sigma, #L4516) and 50 ng/mL interferon gamma (IFN-γ, Peprotech, #300-02) for M1, or supplemented with 20 ng/mL IL-4 (Peprotech, #200-04) and 20 ng/mL IL-13 (Peprotech, #200-13) for M2, and cultured for an additional 2 days.

### CRISPR-Cas9 Gene Editing

For RNP assembly, sgRNAs (GenScript) were reconstituted in Nuclease-Free Duplex Buffer and pre-complexed with Alt-R *S.p.* Cas9 Nuclease V3 (IDT) at a 2:1 molar ratio (6.2 µM sgRNA:3.1 µM Cas9). A non-targeting control, NTC (IDT), containing a sgRNA with no human genomic target, was used as a control for nucleofection. sgRNA sequences are listed in Supplementary Table S1. For each reaction, sgRNA and Cas9 were mixed in P3 nucleofection solution (Lonza) to a 5 µL volume RNP mix and incubated for ≥ 20 min. Freshly isolated monocytes or HeLa S3 cells were resuspended at 1×10⁶ cells per condition in 15 µL P3. The cell suspension (15 µL) was combined with 5 µL RNP mix, transferred to a 4D-Nucleofector® X cassette (Lonza), and nucleofected using program CM-137, EA-100 or DK-100 with P3 buffer for monocytes or program CN-114 with SE buffer for HeLa. Immediately upon nucleofection, cells were collected into 1 mL total of pre-warmed culture medium and used for downstream applications.

### PCR Amplification, Sequencing s Assessment of Gene Editing Efficiency

Three days post nucleofection, DNA was extracted from HeLa S3 cells or monocytes in monoculture using DNeasy Blood & Tissue (Qiagen) and quantified in a NanoDrop OneC (Thermofisher). For each gene editing condition, a region of 300-700 bp surrounding the predicted sgRNA cut site was amplified by PCR using target-specific primers (IDT, Supplementary Table S2) and Platinum SuperFi II DNA Polymerase (Invitrogen). PCR amplification was performed in a SimpliAmp Thermal Cycler (ThermoFisher). PCR products were purified with the GFX PCR DNA and Gel Band Purification Kit (Cytiva) and mixed with one of the primers used in PCR amplification located at least 300 bp away from the cut-site, and Sanger sequenced by STAB VIDA. Chromatograms of each targeted condition were compared against amplified matched NTC-region using established deconvolution algorithms: ICE (Inference of CRISPR Edits, Synthego)^23^, TIDE (Tracking the Indels by Decomposition, NKI)^24^, DECODR (Deconvolution of Complex DNA Repair)^25^, and SeqScreener (ThermoFisher)^26^, for estimation of editing frequencies (indels and frameshift mutations).

### Metabolic Activity Assessment

Metabolic activity was evaluated on monocytes in monoculture for 7 days. Culture medium was removed, and cells were incubated with PrestoBlue diluted 1:10 in culture medium, for 90 min at 37 °C. After incubation, 100 μL of supernatant was transferred to a flat-bottom 96-well Black Plate (Costar). Fluorescence was read at 560 nm excitation with 590 nm emission wavelength on an Infinite® 200 PRO Micro Plate Reader (Tecan Life Sciences).

### Cell Biomass Determination

Adherent cell biomass was evaluated on monocytes in monoculture for 7 days. Culture medium was removed, and cells were washed three times with DPBS containing Ca^2+^ and Mg^2+^ (+/+) to eliminate detached cells. The remaining adherent cells were fixed in 4% paraformaldehyde (PFA), 4% sucrose for 10 min. Cells were stained with 10% Crystal Violet (Merck) in 20% Methanol (Merck), for 20 min, under orbital agitation. Excess dye was washed with DPBS and the plate was left to dry for at least 24 h. Crystal violet crystals were solubilized in 10% Acetic Acid (Merck) under orbital agitation for 20 min. Plate absorbance was read at 592 nm, on an Infinite® 200 PRO Micro Plate Reader (Tecan Life Sciences).

### 3D Co-Culture of Monocytes and BT474 cells

BT474 breast cancer cells and primary monocytes were mixed in a ratio of 1:1.6, with 25,000 tumour cells and 40,000 monocytes *per* well, accounting for BT474 growth and monocyte loss due to stress induced by the KO procedure. Cells were plated in 96-well Ultra-Low Attachment (ULA) plates (Costar) and centrifuged at 300 x g for 5 min to induce multicellular spheroid formation. Co-cultures were kept in RPMI 1640 without Phenol Red (Gibco), supplemented with 10% (v/v) FBS and 1% (v/v) Penicillin-Streptomycin (P/S), and maintained in a humidified incubator at 37 °C, 5% CO2, for 7 days, with half of the medium replaced twice a week.

### Flow Cytometry Assessment of Polarization Markers s Lectin Expression

Tumor-macrophage co-cultures were collected at day 7 of co-culture and dissociated in TrypLE Express (Gibco) for 15 min at 37°C under agitation. Cells remaining attached to the wells were detached using TrypLE Express and subsequently combined with the dissociated spheroid fraction. Cells were incubated with Human TruStain FcX Fc receptor blocking solution (1:20; BioLegend, #422302) for 10 min at 4°C, followed by staining with conjugated antibody (Supplementary Table S3) for 60 min at 4°C. Dead cells were stained with Ghost Dye Violet 510 Fixable Viability Dye (1:1000; Cell Signaling Technology, #59863) for 15 min at 4°C. Cells were resuspended in FACS buffer (containing Ca²⁺ and Mg²⁺, 2% FBS) and acquired using a Sony ID7000 spectral flow cytometer. Spectral unmixing and initial analysis were performed using ID7000 software, with subsequent analysis of exported FCS files in FlowJo (v10.10.1, BD Biosciences).

### Immunofluorescence Assessment of TAM Infiltration

Spheroids were collected at day 7 of co-culture, washed with DPBS containing Ca²⁺ and Mg²⁺, and fixed with 4% PFA, 4% sucrose for 20 min. Spheroids were permeabilized and blocked for 30 min in DPBS containing Ca²⁺ and Mg²⁺, 2% Triton X-100 (Sigma-Aldrich), 0.2% fish skin gelatin (FSG, Sigma-Aldrich). Spheroids were incubated with an Alexa Fluor 488-conjugated anti-human CD45 antibody (1:50; BioLegend, #304019) in DPBS containing Ca²⁺ and Mg²⁺, 0.2% FSG for 2 h at room temperature followed by overnight incubation at 4°C. Nuclei were stained with 4′,6-diamidino-2-phenylindole (DAPI, 1:1000, ThermoFisher, #D3571) for 20 min at room temperature. Samples were mounted using RapiClear 1.49 (SunJin Lab). Images were acquired using a MICA imaging system in confocal mode (Leica Microsystems) and processed in Fiji/ImageJ.

### TAM Sorting and RNA Extraction

BT474 and macrophage spheroids were collected, dissociated and stained as described for flow cytometry. Viable CD45⁺ cells were isolated by fluorescence-activated cell sorting based on CD45-BV605 staining using a MA900 cell sorter (SONY) with a 100 µm sorting chip in semi-purity mode. Sorted cells were collected directly into RLT buffer (Qiagen) supplemented with 10% v/v β-mercaptoethanol (Sigma), snap-frozen in liquid nitrogen, and stored at −80°C. RNA was extracted using the RNeasy Mini Kit (Qiagen, #74106) with on-column DNase digestion for 15 min. RNA concentration was assessed by NanoDrop, and RNA integrity evaluated using a RNA ScreenTape with a TapeStation 4200 system (Agilent). Samples with RNA integrity number equivalent (RINe) ≥ 8 were used for bulk RNA sequencing.

### Bulk RNA Sequencing of Sorted TAMs

Library preparation, sequencing, and gene-level quantification were performed by Novogene. Briefly, mRNA was enriched using poly-T oligo-attached magnetic beads, libraries were prepared following fragmentation and cDNA synthesis, and sequencing was performed on an Illumina platform. Raw reads were processed with *fastp*, aligned to the reference genome using *HISAT2*, and gene-level counts were generated with *featureCounts*. Raw count matrices were used for downstream analysis.

### Analysis of TAM RNA Sequencing Data

Raw gene counts were analyzed in R using DESeq2 with a paired design including donor and treatment condition *(∼ donor + condition*), comparing lectin-targeted or no electroporated (NO EP) conditions against non-targeting control (NTC). Genes with counts ≥ 10 in at least four samples were retained. Nominal p-value < 0.001 was used as an exploratory threshold for differential expression. Variance-stabilized counts were used for Principal Component Analysis (PCA). For visualization of treatment-associated effects, donor variation was removed using *limma::removeBatchEffect*, while preserving the treatment design. GSEA was performed with *clusterProfiler* using Hallmark gene sets from *msigdbr*, with genes ranked by the DESeq2 Wald statistic. Gene sets with Benjamini–Hochberg adjusted p-value < 0.05 were considered significant.

### Bioinformatics s Computational Analysis

All RNA-sequencing analyses were performed in R (v4.2.2) within the RStudio environment (v2025.5.0.496), using base R functions and the *tidyverse* package suite (v2.0.0; including *dplyr* v1.1.4, *tidyr* v1.3.1, *ggplot2* v3.5.1).

#### Principal Component Analysis

PCA was performed on variance-stabilized expression values generated using *DESeq2::vst*. Global PCA was used to assess the major sources of transcriptional variation across samples. For exploratory visualization of treatment-associated variation, the donor-associated variation was removed using *limma::removeBatchEffect*, while preserving experimental condition in the design. Pairwise PCAs comparing each lectin-targeted condition with NTC were subsequently performed using the 500 most variable genes.

#### Differential Gene Expression Analysis (DGEA)

Raw gene counts were analyzed using DESeq2 with a paired design incorporating donor and experimental condition *(∼ donor + condition)*. Genes with ≥ 10 counts in at least four samples were retained for analysis. NTC was compared with NoEP condition to assess effects associated with RNP delivery, while lectin-targeted conditions were compared with NTC, to assess the effects of lectin perturbation. For exploratory gene-level visualization, genes with nominal p-value < 0.001 and |log₂ fold change| ≥ 1 were highlighted in volcano plots.

#### Gene Set Enrichment Analysis (GSEA)

Gene Set Enrichment Analysis (GSEA) was performed using *clusterProfiler::GSEA* against the Hallmark gene-set collection obtained from *msigdbr*. Genes were ranked according to the DESeq2 Wald statistic, with positive normalized enrichment scores (NES) indicating enrichment in the lectin-targeted condition and negative values indicating enrichment in NTC. Gene sets with Benjamini–Hochberg false discovery rate (FDR) < 0.05 were considered significantly enriched. For the combined Hallmark summary, pathways were displayed when they represented either a recurrent response, defined as FDR < 0.05 in at least two of the five lectin perturbations, or a strong target-specific response, defined as FDR < 0.05 together with |NES| ≥ the 90th percentile across all lectin-targeted vs NTC comparisons.

#### PROGENy Pathway Activity Analysis

Signaling-pathway activity was inferred using *PROGENy* through the *decoupleR* package. The human PROGENy model comprising the top 500 responsive genes per pathway was applied to TMM-normalized, voom-transformed expression values using the multivariate linear model implemented in *decoupleR::run_mlm*. Differences in inferred pathway activity were subsequently tested using *limma* with a paired design incorporating donor and condition *(∼ donor + condition)*, and p-values were corrected for multiple testing using the Benjamini–Hochberg method.

## Results

### Benchmarking Nucleofection Programs for CRISPR–Cas9 RNP Editing in Primary Human Monocytes

To establish and benchmark a CRISPR–Cas9 RNP editing workflow in primary human monocytes (Figure 1A), *CD14*, a highly expressed monocyte surface co-receptor that provides a readily quantifiable flow-cytometric readout of target depletion, was selected as a positive-control target (Figure 1B).^17^ Three nucleofection programs were selected for comparison: CM-137 and DK-100, previously used for CRISPR–Cas9 RNP editing in primary human monocytes, and the equipment manufacturer-recommended EA-100 program.^17,18,27^

**Figure 1.**
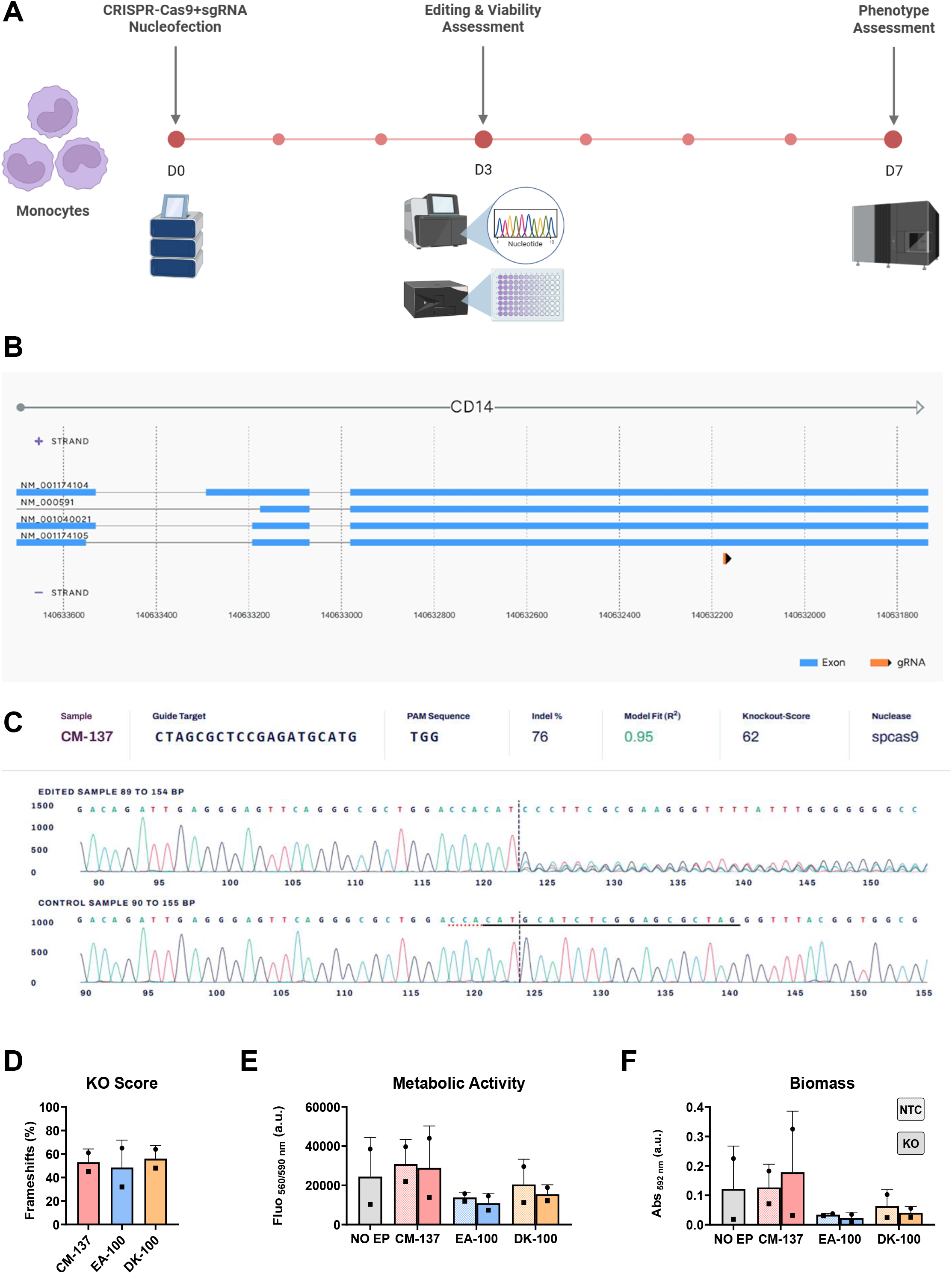
Qualification of CRISPR–Cas9 ribonucleoproteins delivery and editing assessment in primary human monocytes. A. Schematic of the experimental workflow. Primary human monocytes were nucleofected with pre-assembled non-targeting control or CD14-targeting Cas9–sgRNA RNPs using CM-137, EA-100, or DK-100 nucleofection programs, and assessed for gene editing efficiency, cell compatibility and phenotypic stimuli response. B. Schematic representation from GenScript of the CD14 genomic locus showing annotated RefSeq transcript isoforms, exon organization (blue), and the position of the CD14-targeting sgRNA (orange). C. Representative Sanger-trace chromatograms for matched non-targeting control (NTC) and CD14-targeting conditions, deconvoluted using ICE (Inference of CRISPR Edits, Synthego)^23^ in monocytes nucleofected using CM-137. The sgRNA sequence (black line), protospacer adjacent motif (PAM, dotted red line), predicted Cas9 cleavage site (dashed black line), indel percentage, predicted knockout score, and model fit are indicated. D. ICE-derived knockout score based on predicted frameshift mutations following CD14 editing using CM-137, EA-100, or DK-100. E. Metabolic activity assessed by resazurin reduction. F. Retained adherent biomass assessed by crystal violet staining. No EP denotes non-electroporated cells. Data are from two independent healthy donors, represented as mean ± SD.

Monocytes from two independent healthy donors were nucleofected with CD14-targeting or NTC pre-complexed RNPs, and editing was evaluated by Sanger sequencing. Chromatograms were analyzed using four deconvolution algorithms: ICE (Inference of CRISPR Edits, Synthego)^23^, TIDE (Tracking the Indels by Decomposition, NKI)^24^, DECODR (Deconvolution of Complex DNA Repair)^25^, and SeqScreener (ThermoFisher)^26^. Estimated editing frequencies (indels and frameshift mutations) were broadly concordant between algorithms, while ICE consistently yielded the highest model fit across the three nucleofection programs, reflecting the agreement between the observed Sanger chromatogram and the sequence mixture predicted by the deconvolution model (Supplementary Figure S1). ICE was therefore selected as the primary method for subsequent editing analyses.

Matched NTC and CD14-targeting Sanger chromatograms showed concordant traces upstream of the predicted Cas9 cleavage site, followed by mixed-base calls in the CD14-targeted sample downstream of the expected cleavage site (Figure 1C). All three nucleofection programs supported substantial and comparable *CD14* editing according to estimated indels (Supplementary Figure S1) and predicted frameshift mutations (Figure 1D), despite donor-dependent variability in efficiency.

Cellular compatibility was therefore evaluated alongside genomic editing. Across the two donors, neither resazurin reduction nor crystal violet measurements showed a consistent difference between CD14-targeted and matched NTC conditions within each nucleofection program (Figure 1E-F). Among the programs tested, CM-137 showed the greatest overall retention of metabolic activity and adherent cellular biomass, with values approaching those observed in non-electroporated controls. Given its comparable editing efficiency and superior cellular compatibility, CM-137 was selected for subsequent experiments.

Additional pilot experiments examined whether the Alt-R Cas9 Electroporation Enhancer could improve CD14 editing, as previously reported.^17^ Although CD14 targeting substantially reduced CD14 expression at day 7, we could not independently confirm an additional benefit of enhancer supplementation under our experimental conditions (Supplementary Figure S2A). Additionally, representative microscopy images further suggested reduced adherent cell density and altered cell morphology, including increased cell rounding, in enhancer-treated cultures, consistent with reduced viability. Therefore, the electroporation enhancer was not included in subsequent experiments (Supplementary Figure S2B).

To assess whether edited monocytes retained responsiveness to distinct macrophage-polarizing stimuli, CD14-targeted cells were exposed to M1/pro-inflammatory (IFN-γ + LPS) or M2/alternative-polarizing conditions. Despite sustained CD14 depletion, edited cells retained the expected polarization-associated expression patterns, with increased CD80 and reduced CD206 under M1-like conditions, and increased CD206 with lower CD80 expression under M2-like conditions (Supplementary Figure S2C).

Notably, CD80 expression following IFN-γ + LPS stimulation was reduced in CD14-targeted macrophages compared with the corresponding non-edited control. Since CD14 facilitates LPS–TLR4 signaling, with downstream NF-κB activation contributing to the induction of pro-inflammatory co-stimulatory molecules such as CD80, this observation was consistent with attenuated CD80 expression following CD14 depletion.^28^ Owing to the exploratory single-donor design, these data were interpreted qualitatively and were not used to infer a definitive effect of CD14 disruption on macrophage polarization.

Together, these experiments supported a CRISPR–Cas9 RNP workflow in primary human monocytes that enabled efficient editing while preserving cell recovery and responsiveness to macrophage-polarizing stimuli.

### Screening of sgRNAs Targeting Lectins in HeLa S3 Cells

Efficient perturbation of candidate lectin targets first required the identification of active sgRNAs for each gene. Optimizing and selecting efficient sgRNAs directly in primary monocytes is technically demanding and time-consuming, as it requires repeated isolation of primary cells from donors for every guide tested. To circumvent this constraint, sgRNA activity was instead first screened in HeLa S3 cells, a readily available and standardized system that allowed multiple guides to be compared efficiently before their application in primary monocytes (Figure 2A).^29^ Multiple sgRNAs targeting different exonic regions of *CLEC4E*, *CLECcA*, *CD20S*, *SELL*, and *SIGLEC9* were individually evaluated, while the previously used CD14 sgRNA was included as a technical reference control (Figure 2B).

**Figure 2.**
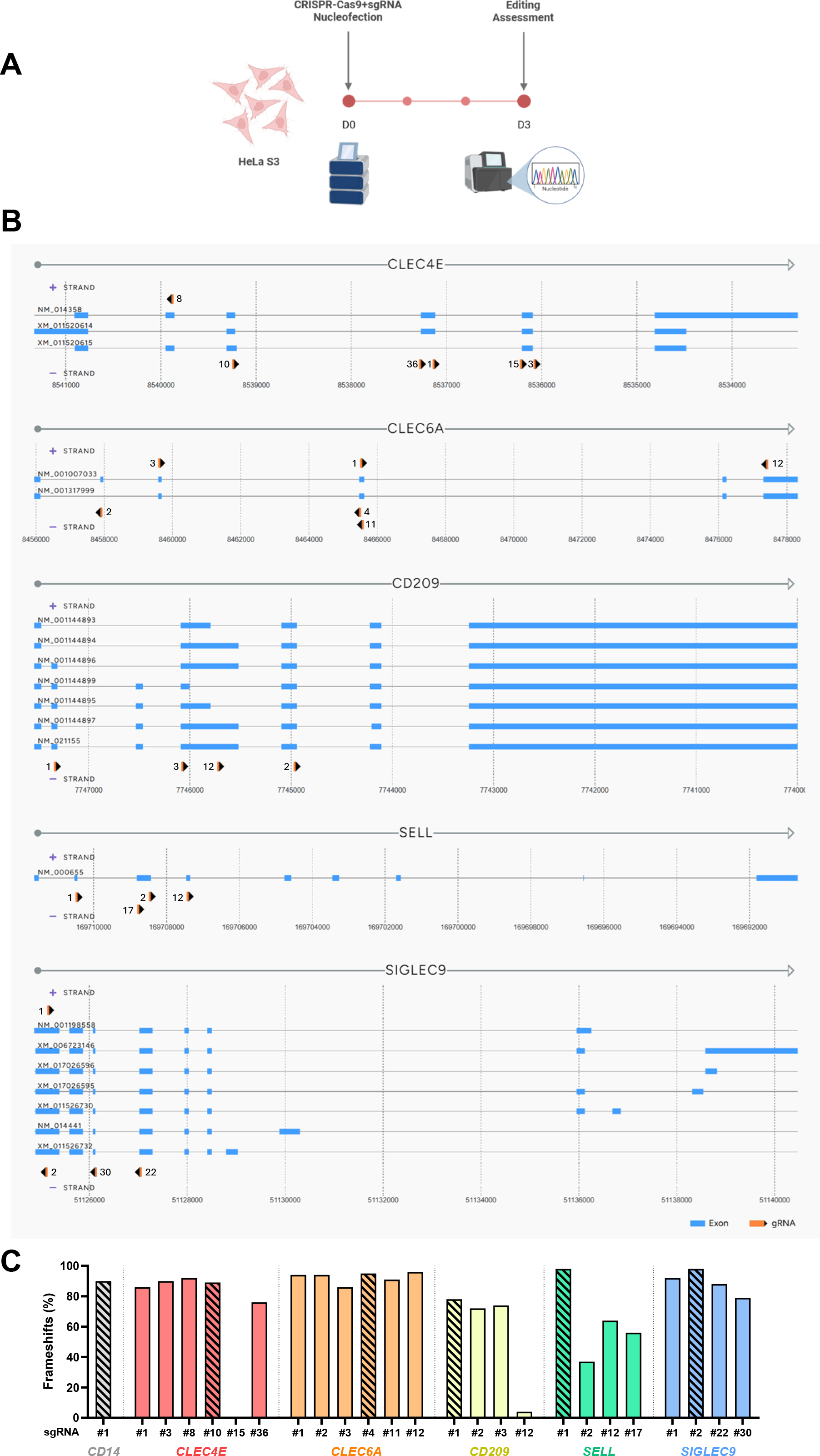
Screening and selection of sgRNAs targeting candidate lectins in HeLa S3 cells. A. Schematic of the sgRNA screening workflow. HeLa S3 cells were nucleofected with pre-assembled Cas9–sgRNA RNPs targeting CLEC4E, CLEC6A, CD209, SELL, or SIGLEC9, with CD14 included as a technical reference control. Genomic editing was assessed three days after nucleofection by Sanger sequencing and ICE analysis to support selection of sgRNAs for subsequent evaluation in primary human monocytes. B. Schematic representation of the lectins genomic locus showing annotated RefSeq transcript isoforms, exon organization (blue), and the position of the targeting sgRNAs (orange). C. Predicted frameshift frequencies determined by ICE analysis following individual sgRNA delivery in HeLa S3 cells. Hatched bars indicate the sgRNA selected for each target for subsequent evaluation in primary human monocytes. Data represent a single technical screening experiment.

HeLa S3 cells were nucleofected with individual Cas9–sgRNA RNPs, and genomic editing was assessed three days later by Sanger sequencing and ICE analysis. Predicted frameshift frequencies varied between sgRNAs within each target, demonstrating the importance of guide-specific screening prior to primary-cell application (Figure 2C). Several guides produced high predicted frameshift frequencies, whereas others showed substantially lower activity. Indel frequencies and ICE model-fit parameters were additionally considered to support interpretation of the screening results (Supplementary Figure S3).

Guide selection prioritized sgRNAs combining higher editing efficiency with targeting of exonic regions shared across the largest number of annotated transcript isoforms. Based on these criteria, CLEC4E sgRNA #10, CLEC6A sgRNA #4, CD209 sgRNA #1, SELL sgRNA #1, and SIGLEC9 sgRNA #2 were selected for subsequent experiments in primary human monocytes (Figure 2C, hatched bars).

As this screen was intended to compare sgRNA performance rather than lectin biology, differences in editing efficiency between targets were not interpreted mechanistically, and the selected guides were next evaluated in the primary-monocyte system.

### BT474–Monocyte Co-Culture Supports Tumor-Associated Macrophage Differentiation and Expression of Target Lectins

Before applying the selected CRISPR–Cas9 RNP workflow to the candidate lectins, the BT474–monocyte co-culture used for perturbation studies was characterized to assess macrophage phenotype, cellular distribution, and expression of the selected targets. Primary human monocytes were co-cultured with BT474 tumor cells in ultra-low attachment plates for seven days, and the CD45⁺ population was analyzed by flow cytometry (Figure 3A).

**Figure 3.**
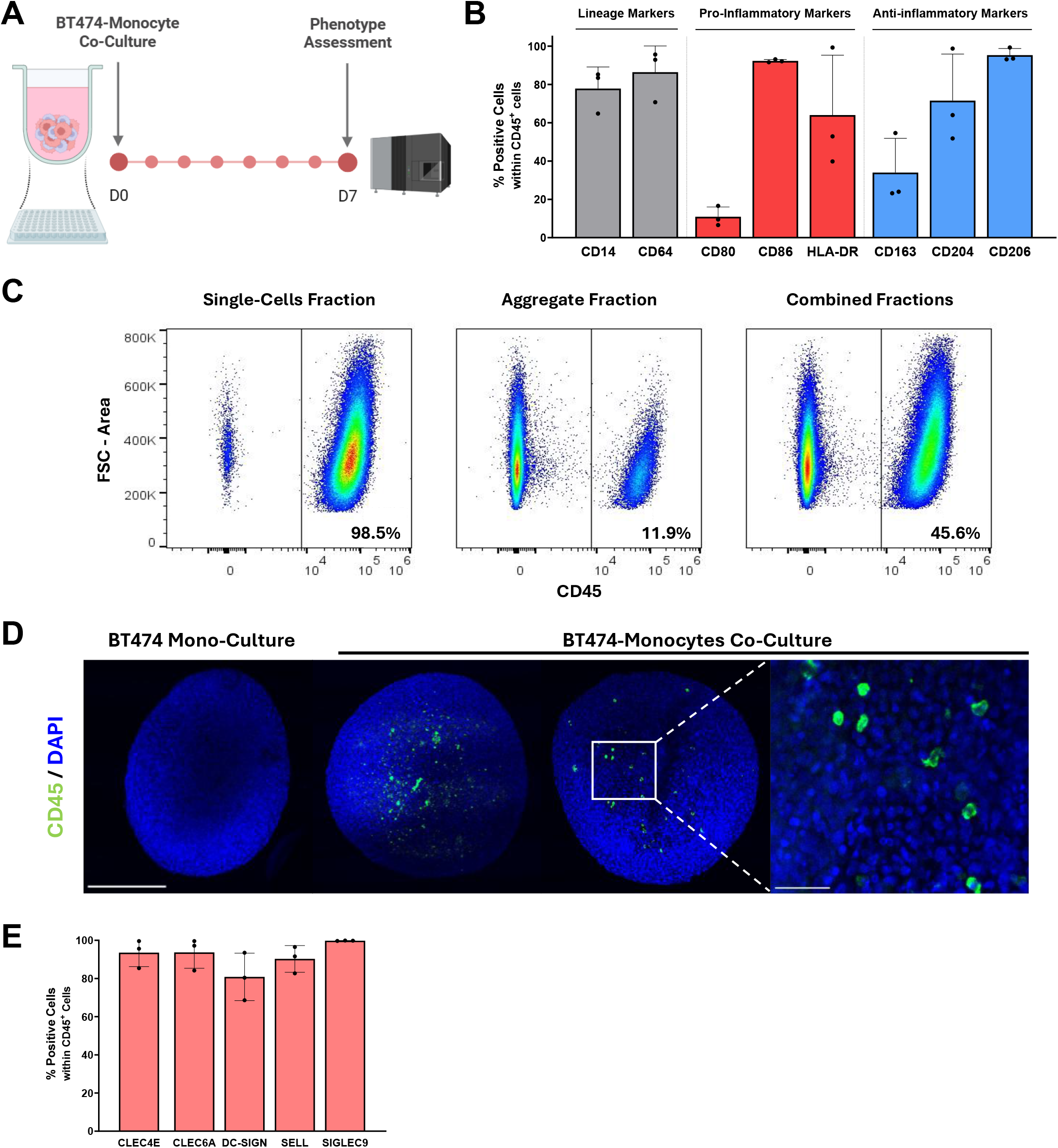
Phenotypic characterization of the BT474–monocyte co-culture used for lectin perturbation studies. A. Schematic representation of the experimental workflow. Primary human monocytes were co-cultured with BT474 spheroids in ultra-low attachment plates for seven days, followed by phenotypic characterization by flow cytometry and immunofluorescence. B. Flow-cytometry analysis of macrophage lineage (CD14 and CD64) and polarization-associated markers at day 7, including CD80, CD86 and HLA-DR (pro-inflammatory-associated markers), and CD163, CD204 and CD206 (anti-inflammatory-associated markers). Data represented as mean ± SD of three independent monocyte donors. C. Representative flow-cytometry analysis of CD45⁺ cell distribution following fractioning of the day-7 co-culture into single-cells, aggregate-associated and combined cell fractions. Percentages indicate the frequency of CD45⁺ cells within each fraction. D. Representative immunofluorescence images showing CD45⁺ cells (green) within BT474 co-culture spheroids. The tumor-cell monoculture was included as a negative control for CD45 staining. Nuclei were counterstained with DAPI (blue). Images were acquired as confocal Z-stacks and are shown as maximum-intensity projections of X optical Z-slices of 2.5 μm thickness for whole-spheroid images or X optical Z-slices of 1 μm thickness for magnified region. Scale bar = 250 μm for whole-spheroid images and 50 μm for the magnified region. E. Flow-cytometry analysis of CLEC4E, CLEC6A, CD209, SELL and SIGLEC9 expression within the CD45⁺ population at day 7. Data represented as mean ± SD of three independent monocyte donors.

At day 7 of co-culture, the CD45⁺ population showed high expression of the myeloid lineage markers CD14 and CD64 (Figure 3B). Co-cultured macrophages additionally displayed a heterogeneous polarization-associated phenotype, with expression of pro-inflammatory markers CD86 and HLA-DR together with anti-inflammatory markers CD163, CD204 and CD206 (Figure 3B). These data supported macrophage differentiation within the co-culture while consistent with polarization into a TAM phenotype.

To assess the distribution and infiltration of monocyte-derived cells within the co-culture spheroid, the different cellular fractions (aggregate and suspension cells) were analyzed separately. CD45⁺ cells accounted for 11.9% of the cells recovered within the aggregate fraction (infiltrated macrophages), whereas the non-aggregate single-cell fraction was predominantly CD45⁺ (98.5%, suspension macrophages); analysis of the combined cell population resulted in an overall CD45⁺ frequency of 45.6% (Figure 3C). Immunofluorescence imaging further confirmed the presence of CD45⁺ cells within co-culture spheroids (Figure 3D). Based on this distribution, aggregate-associated and non-aggregate macrophage populations were subsequently recovered and pooled for downstream analyses. This approach was intended to capture macrophage responses to both direct tumor-cell interactions and soluble tumor-derived signals, including cytokines and secreted or shed glycan-bearing molecules.

Importantly, all five candidate lectins were detectable in CD45⁺ cells at day 7, with no difference in the proportion of positive cells between targets (Figure 3E). Together, these results confirmed expression of the selected targets within the experimental co-culture and supported their subsequent genetic perturbation.

### CRISPR–Cas9 RNP Editing of Candidate Lectins in Primary Human Monocytes

Having established the nucleofection conditions, selected active sgRNAs and confirmed expression of the candidate lectins within the co-culture, the two components of the workflow were combined to evaluate target-specific editing in primary human monocytes. Freshly isolated monocytes were nucleofected using the CM-137 program with the selected lectin-targeting sgRNAs pre-complexed with Cas9 into RNPs and then divided into parallel experimental arms (Figure 4A). Part of the edited cell population was maintained in monoculture for genomic editing assessment at day 3, while remaining cells were incorporated into BT474–monocyte co-cultures and recovered by CD45⁺ cell sorting at day 7 for downstream transcriptomic analysis.

**Figure 4.**
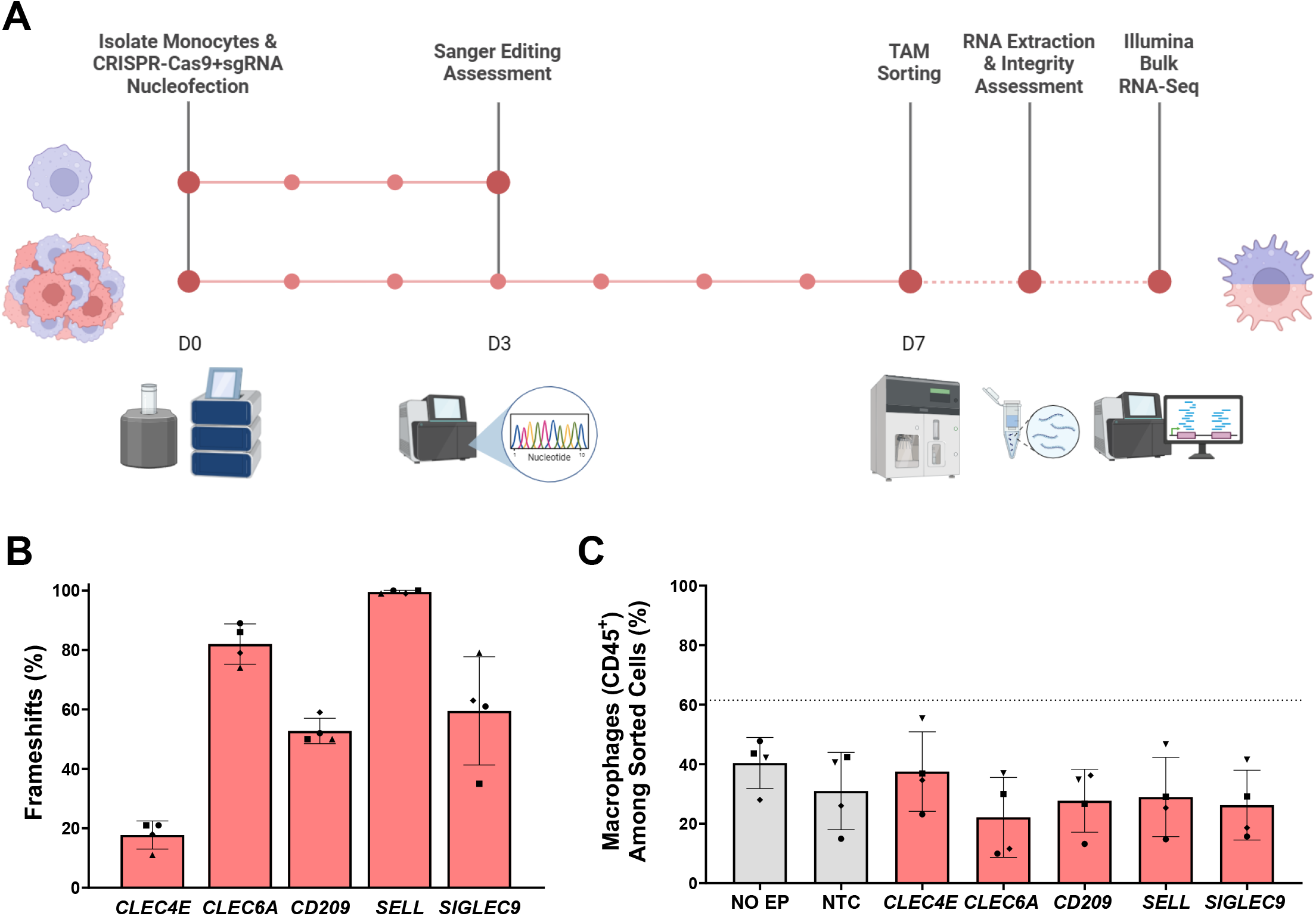
Target-dependent CRISPR–Cas9 editing and macrophage recovery following lectin perturbation. A. Schematic representation of the experimental workflow. Freshly isolated primary human monocytes were nucleofected with CRISPR–Cas9 RNPs at day 0. A parallel aliquot was maintained in monoculture for genomic editing assessment by Sanger sequencing and ICE analysis at day 3, while matched edited cells were incorporated into BT474–monocyte co-cultures. At day 7, CD45⁺ macrophages were isolated by fluorescence-activated cell sorting (FACS), followed by RNA extraction, integrity assessment and bulk RNA sequencing. B. Predicted frameshift frequencies determined by ICE analysis in parallel monoculture samples for CLEC4E, CLEC6A, CD209, SELL, and SIGLEC9. C. Frequency of CD45⁺ macrophages recovered from BT474–monocyte co-cultures at day 7 across non-electroporated (No EP), non-targeting control (NTC), and lectin-targeting conditions. Data represented as mean ± SD of four independent healthy donors.

ICE analysis revealed substantial target-dependent differences in predicted frameshift mutations (Figure 4B), indicating that the magnitude of sgRNA activity observed in HeLa S3 cells did not translate uniformly to primary human monocytes, expectedly due to the influence of cellular context on CRISPR–Cas9 editing efficiency. *CLEC4E* showed consistently lower predicted disruption (around 20%), whereas *CLEC6A* and *SELL* reached high to near-complete disruption in all donors (above 80%). *CD209* displayed intermediate and relatively consistent disruption, while *SIGLEC9* exhibited intermediate-to-high editing, with greater inter-donor variability. Total indel frequencies followed a similar target-dependent pattern, with consistently high ICE model-fit values supporting the corresponding editing estimates (Supplementary Figure S4A–B).

The proportion of CD45⁺ macrophages recovered from the co-cultures varied between donors (Figure 4C). However, no lectin-targeted condition consistently reduced the macrophage fraction relative to NTC, supporting comparable macrophage recovery for downstream analysis. Additionally, representative backsort analysis confirmed 99.8% purity of the isolated CD45⁺ population (Supplementary Figure S4C), and RNA extracted from the sorted macrophages showed consistently high integrity (RINe) across control and lectin-targeted conditions (Supplementary Figure S4D).

Together, these results demonstrated that the established workflow enabled target- and donor-dependent genetic disruption of the five candidate lectins while maintaining macrophage populations suitable for subsequent transcriptomic analysis.

### Lectin Perturbation Induces Target-Dependent Transcriptional Remodeling in Tumor-Associated Macrophages

Bulk RNA sequencing was performed on recovered CD45⁺ macrophages across NoEP, NTC, and the five lectin-targeting conditions. Principal component analysis of global expression profiles revealed a pronounced donor-dependent structure, with samples clustering predominantly according to donor rather than experimental condition (Figure 5A). Accordingly, donor identity was incorporated into the differential-expression model to account for the paired experimental design. Following removal of the donor-associated component for exploratory visualization, pairwise PCA revealed condition-associated separation between NTC and each lectin-targeted condition, although partial overlap remained for *CLEC6A* (Supplementary Figure S5A–E).

**Figure 5.**
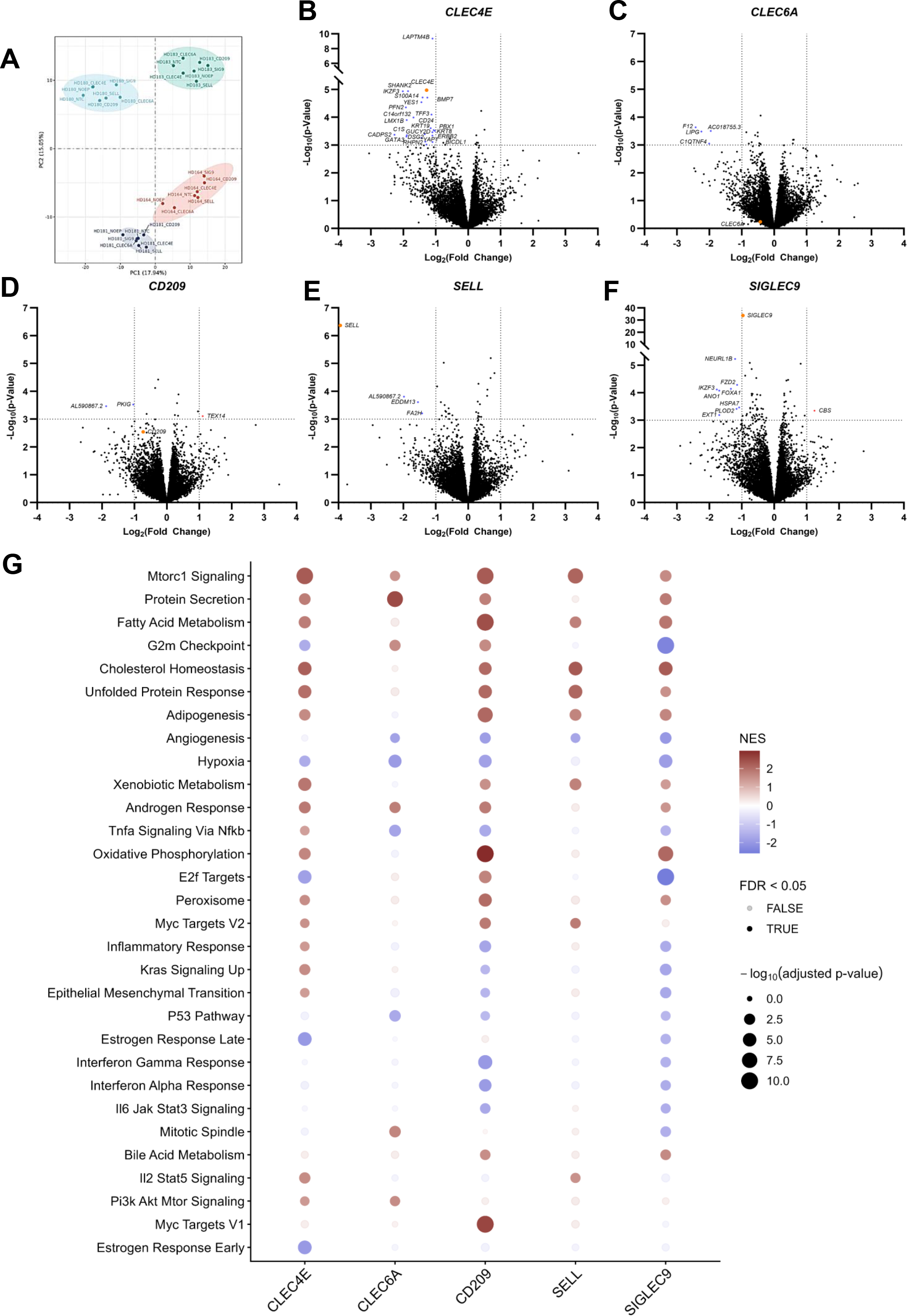
Transcriptomic remodeling of co-cultured macrophages following lectin perturbation. A. Principal component analysis of RNA-seq profiles from four donors across no electroporation control (NoEP), non-targeting control (NTC) and the five lectin-targeting conditions. Samples are colored by donor, with shaded ellipses indicating donor-associated clustering. B–F. Volcano plots showing exploratory gene-level changes in B, CLEC4E; C, CLEC6A; D, CD209; E, SELL; and F, SIGLEC9-targeted macrophages relative to NTC. Dotted lines indicate ∣log2FC∣=1 and nominal p=0.001; genes meeting both thresholds or targeted by sgRNA are highlighted and labelled. G. Hallmark gene set enrichment analysis (GSEA) across the five lectin-targeting conditions relative to NTC. Dot color represents normalized enrichment score (NES), with positive values indicating enrichment in the targeted condition and negative values enrichment in NTC; dot size represents −log10 FDR-adjusted p-value and opacity indicates FDR < 0.05. Displayed pathways were recurrently significant across lectin perturbations or showed a strong target-specific enrichment.

The transcriptional impact of RNP delivery itself was first examined by comparing NTC with NoEP macrophages. Exploratory gene-level analysis indicated only minimal transcriptional differences between these controls (Supplementary Figure S5F), supporting the use of NTC as the matched reference for subsequent lectin-targeting comparisons.

At the individual gene level, the extent of transcriptional change varied across lectin-targeted conditions (Figure 5B–F). *CLEC4E* targeting showed the strongest response, with 24 downregulated genes, followed by *SIGLEC9*, with nine downregulated and one upregulated gene. More limited transcriptomic changes were observed following *CLEC6A* and *SELL* targeting, each with four downregulated genes, whereas *CD209* showed the most modest gene regulation. Although the number of genes meeting significance varied substantially between targets, coordinated biological responses may still emerge from more subtle, concordant transcriptional changes across larger gene sets, so ranked gene-set enrichment analysis was used to assess pathway-level responses.

Hallmark GSEA revealed broader coordinated transcriptional remodeling following lectin perturbation (Figure 5G). The enriched programs were target-dependent but recurrently involved lipid and other metabolic processes. mTORC1 signaling was positively enriched across all five perturbations, while cholesterol homeostasis, fatty-acid metabolism and oxidative phosphorylation were recurrently enriched together with related biosynthetic and stress-response pathways. In contrast, hypoxia- and angiogenesis-associated programs showed relative depletion following several perturbations, while inflammatory and cytokine-associated pathways were modulated in a target-dependent manner.

The combination of pathways affected differed markedly between individual lectins. Despite the relatively limited gene-level changes observed for *CLEC6A*, its perturbation produced coordinated pathway-level remodeling, with enrichment of mTORC1/PI3K–AKT– mTOR and protein-secretion programs accompanied by depletion of hypoxia-, angiogenesis- and TNFα/NF-κB-associated signatures. *CD20S* and *SIGLECS* showed broader and partially overlapping responses, characterized by enrichment of lipid and oxidative metabolic programs, together with reduced hypoxia-, angiogenesis- and inflammatory-associated signatures. *SELL* targeting showed a comparatively more restricted response, predominantly involving metabolic and secretory programs. Notably, *CLEC4E* targeting was associated with extensive lipid, oxidative, mTORC1 and stress-response remodeling despite its comparatively low editing efficiency.

As a complementary approach, PROGENy analysis was performed to examine whether the transcriptional patterns identified by Hallmark GSEA were reflected at the level of inferred signaling-pathway activity. The analysis revealed target-dependent modulation across several signaling pathways (Supplementary Figure S5G). Reduced hypoxia-associated activity was most pronounced following *CD20S* and *SIGLECS* perturbation, consistent with the depletion of hypoxia-related transcriptional programs observed by Hallmark GSEA. Increased inferred PI3K activity was also observed across several lectin-targeting conditions, including *CLEC6A*, complementing the enrichment of PI3K–AKT– mTOR/mTORC1-associated programs identified in GSEA. Overall, PROGENy analysis provided a complementary signaling-level view of the target-dependent transcriptional remodeling identified by gene-set enrichment analysis.

Collectively, these results demonstrate that perturbation of the selected lectins modifies the transcriptional state of macrophages within the tumor co-culture. While the specific response differed between targets, recurrent modulation of lipid-associated and oxidative metabolic programs indicates a shared impact on macrophage state regulation.

## Discussion

This study established a CRISPR–Cas9 RNP workflow for comparative perturbation of candidate myeloid lectins in primary human monocytes and applied it to five receptors during tumor-conditioned macrophage differentiation. Despite target- and donor-dependent editing efficiencies, perturbation of the selected lectins consistently affected coordinated transcriptional programs, supporting a functional contribution to TAM-state regulation.

The transcriptomic analysis suggests that perturbation of the selected lectins modifies the TAM state predominantly at the level of coordinated biological programs rather than through extensive changes in individual transcripts. The most biologically consistent consequence of lectin perturbation was the remodeling of metabolic programs. The recurrence of mTORC1 signaling, cholesterol homeostasis, fatty-acid metabolism and oxidative metabolism across independently targeted receptors suggests these responses extend beyond individual lectin-specific pathways and involve metabolic processes associated with TAM adaptation. This is particularly relevant given the increasing evidence that lipid metabolism is an active determinant of TAM phenotype and function.^30–32^ Lipid accumulation, fatty-acid oxidation, and mitochondrial metabolism have been linked to tumor-supportive macrophage states, while altered cholesterol trafficking can directly regulate TAM polarization and function.^30–33^ More recent work in glioblastoma has further shown that lipid-enriched TAM populations can provide metabolic support to tumor cells and that manipulation of cholesterol and oxysterol metabolism can redirect macrophage signaling and cell fate.^34,35^ These studies reinforce the view that remodeling of TAM lipid metabolism is not merely a consequence of polarization, but part of the machinery that sustains macrophage adaptation within tumors.

Accordingly, PI3K–AKT–mTORC1 signaling can regulate metabolic adaptation and functional state in macrophages, with its effects on tumor-associated myeloid phenotypes strongly shaped by the surrounding tumor context.^36^ This raises the hypothesis that structurally distinct lectin receptors converge on some shared immunometabolic nodes, while exerting different effects on inflammatory, hypoxic and angiogenic programs. Such interactions between macrophage metabolism and angiogenic/hypoxic regulation have previously been demonstrated experimentally in tumors.^37^ Direct lipid, cholesterol/oxysterol and central carbon metabolic flux profiling of lectin-perturbed macrophages would therefore be particularly informative in determining whether these transcriptional changes correspond to altered lipid uptake, storage, synthesis, or utilization and how such changes contribute to modulation of immunosuppressive TAM features.

Several lectin perturbations attenuated hypoxic and angiogenic signatures, but inflammatory, TNF/NF-κB and interferon-associated programs were also reduced in selected conditions. In tumors, sustained inflammatory and NF-κB signaling can promote angiogenesis, tissue remodeling, tumor-cell survival and immune dysfunction, linking these inflammatory programs to tumor-supportive TAM functions.^38^ This distinction is relevant to lectin biology as tumor-associated glycan recognition can simultaneously induce inflammatory mediators and tumor-supportive functions. ^8^

The individual lectins nevertheless provide distinct biological perspectives. The response to *CLEC6A* (Dectin-2) targeting combined coordinated pathway-level remodeling, including attenuation of TNF/NF-κB-, hypoxia- and angiogenesis-associated programs despite limited individual-gene changes. The accompanying reduction of endothelial lipase (*LIPG*) provides a gene-level connection to this phenotype, given the established role of this enzyme in regulating macrophage lipid composition and inflammatory cytokine production.^39^ This is noteworthy given the established role of Dectin-2 as an activating C-type lectin receptor, while its engagement has also been implicated in macrophage-mediated antitumor activity and explored as a TAM-directed agonistic strategy.^12,40,41^ The contrast raises the possibility that Dectin-2 activity in tumor-conditioned macrophages is likely highly shaped by ligand availability, signaling strength and the pre-existing macrophage state.

*CLEC4E* (Mincle) targeting was technically more challenging but potentially equally biologically important. Despite limited editing efficiency, a coherent metabolic and mTORC1-associated response was nevertheless observed. This is particularly relevant given emerging evidence that tumor-derived lipids can engage Mincle and promote lipid accumulation and pro-tumorigenic macrophage activity, together with independent studies implicating Mincle in macrophage-dependent tumor immune suppression.^10,11,42^ Importantly, antibody blockade or genetic deficiency of Mincle has been shown to reduce lipid accumulation, ER-stress-associated signaling and suppressive activity in tumor-conditioned macrophages, while macrophage-specific Clec4e deletion more recently altered TAM state and enhanced antitumor immune features.^10,11^ The present findings therefore raise the hypothesis that metabolic regulation may represent one mechanism of Mincle-dependent TAM regulation, and its inefficient disruption should consequently be viewed as a rationale to warrant improved genetic interrogation.

The responses observed for *CD209* (DC-SIGN) and *SIGLEC9* targeting similarly place this work in line with growing interest in tumor glycan–myeloid interactions. Both receptors have established or emerging roles in sensing aberrant tumor glycosylation, including fucosylated ligands for DC-SIGN and sialylated ligands for Siglec-9, with downstream effects on macrophage differentiation and immune regulation.^8,9,43–46^ For *CD20S*, this is particularly relevant in breast cancer, where engagement of DC-SIGN on macrophages by fucosylated tumor-derived ligands has been associated with a proangiogenic macrophage profile.^8,45,46^ The depletion of hypoxia- and angiogenesis-associated programs following *CD209* perturbation is therefore directionally consistent with this reported function, although functional consequences were not directly assessed here. Similarly, cancer-associated sialylated ligands engaging Siglec-9 have been shown to influence monocyte differentiation toward tumor-associated macrophage states, providing a biological context for the transcriptional remodeling observed following *SIGLECS* perturbation.^9,43,44^ Although neither perturbation produced a uniform pro-inflammatory shift in our model, direct blockade of DC-SIGN in bladder cancer TAMs and Siglec-9 in ovarian cancer TAMs has been reported to reduce immunosuppressive macrophage features and enhance CD8⁺ T-cell activity, supporting functional effects that are likely dependent on tumor context.^44,47^ Together, these findings support a contribution of glycan-dependent signaling through DC-SIGN and Siglec-9 to tumor-conditioned macrophage states.

*SELL* targeting, by contrast, produced a more restricted transcriptional response despite efficient disruption. This may reflect the predominant function of L-selectin in leukocyte adhesion and recruitment across the vascular endothelium, processes that are largely absent from the present tumor cell–macrophage co-culture.^13^ Consistent with this interpretation, L-selectin deficiency or functional blockade in tumor models has been shown to impair leukocyte recruitment to sites of tumor-cell arrest and reduce metastatic establishment.^48^ Accordingly, the functional consequences of SELL perturbation may be more apparent in models incorporating endothelial barriers, leukocyte trafficking or tissue recruitment.

These comparative analyses depended on first achieving efficient lectin perturbations across technically challenging primary human monocytes. Regarding the deconvolution algorithm for analysis of Sanger chromatograms, ICE choice was consistent with comparative analysis showing reliable performance between methods for relatively simple, short-indel editing outcomes.^23,49^ As to published workflows for CRISPR–Cas9 RNP delivery to primary human monocytes, these remain limited and employ distinct nucleofection settings. Freund et al.^17^ used the CM-137 program for human monocyte editing following its selection in broader myeloid-cell screens, whereas Hiatt et al.^18^ independently identified DK-100 during optimization in primary human monocytes, and EA-100 is recommended by the manufacturer for nucleofection of primary human monocytes.^27^ However, to our knowledge, these programs had not been directly compared in parallel within the same primary human monocyte CRISPR–Cas9 RNP workflow. Therefore, CM-137, EA-100 and DK-100 were benchmarked under matched experimental conditions. The selection of *CD14* for this comparison, a highly expressed marker in monocytes, provided a readily detectable surface readout for the assessment of target depletion, and supported identification of CM-137 as the most favorable compromise between editing efficiency and cellular compatibility.

The HeLa S3 screen was used as a relative sgRNA-ranking strategy before validation in primary human monocytes. Differences in cellular context, including chromatin accessibility and locus-specific features, were expected to influence the absolute efficiency achieved after transfer to primary cells.^50^ Yet, sgRNAs showing the highest activity within each target in HeLa S3 cells would have a greater likelihood of retaining the best activity in monocytes. While sgRNAs targeting *CLEC6A* and *SELL* retained high activity, disruption of *CLEC4E* was substantially reduced despite its strong performance in the initial screen. Thus, the HeLa S3 screen served as a guide-enrichment strategy, while direct assessment in the primary-cell context remained necessary to establish target-specific editing efficiency. Overall, the workflow enabled perturbation of all five lectin candidate genes while preserving macrophage recovery and providing a suitable platform for subsequent functional interrogation.

The platform developed here provides a direct route to further dissect how the selected lectins contribute to TAM regulation. Combinatorial perturbation studies could determine whether different lectins act independently, cooperatively or with partially overlapping functions in TAM immunometabolic regulation, while paired-sgRNA strategies would be particularly valuable for technically challenging receptors such as *CLEC4E*. Protein-level validation, selected sorting and rescue approaches could further strengthen receptor-specific mechanistic attribution. Additionally, an important open question concerns how lectin perturbation in macrophages affects the co-cultured tumor cells themselves, an aspect not addressed by the present analysis. Extending bulk RNA-seq to the BT474 tumor-cell fraction recovered from the same co-cultures would allow direct assessment of paracrine effects of lectin-perturbed macrophages on tumor-cell transcriptional state, complementing the macrophage-analysis presented here. Applying the workflow to co-culture models with tumor cell lines with distinct glycosylation profiles would help establish how tumor context determines lectin dependency. Most importantly, integration of genetic perturbation with lipidomics, glycomics, and functional assays, including phagocytosis, cytokine secretion, tumor-cell support, angiogenic activity, and macrophage-T-cell interactions, would help establish how the metabolic and transcriptional remodeling identified here translates into modification of immunosuppressive TAM functions.

Overall, these findings support lectin signaling as a functional regulatory layer connecting environmental sensing with the metabolic and transcriptional adaptation of TAMs. Rather than inducing a uniform pro-inflammatory transition, perturbation of individual receptors modified distinct but overlapping components of the TAM state. Within this framework, *CLEC6A* and *CLEC4E* emerge as particularly compelling candidates for deeper mechanistic investigation, adding to an emerging body of evidence linking these receptors to TAM regulation. In parallel, the responses observed following *CD20S* and *SIGLECS* perturbation reinforce existing evidence for tumor glycan–myeloid signaling as a regulator of macrophage state. These results therefore identify lectin-dependent immunometabolic regulation as a promising axis for future modulation of immunosuppressive TAM states.

## Acknowledgments

The authors thank Olivia Rodrigues and the iBET Flow Cytometry Unit for technical expertise, support with TAM sorting, and access to state-of-the-art instrumentation, and Inês A. Isidro (iBET & ITQB NOVA) for her feedback during manuscript preparation.

## Additional Elements

### Ethics Approval and Consent to Participate

The use of anonymized human buffy coats of healthy blood donors from the Portuguese Institute for Blood and Transplantation (Instituto Português do Sangue e da Transplantação, IPST, Lisbon, Portugal) was approved by the ethical review board of IHMT – ITQB – NSL – IGC for the project “Descodificar interações glícido-lectina típicas de macrófagos imunossupressores em Cancro - GLYCO-TAM” (reference “Parecer 16.23”). Written informed consent for blood donation and the research use of the donated material was obtained from all donors by the IPST before sample collection. Buffy coats corresponded to blood donations collected by the IPST on the day preceding their provision to the investigators. Samples used in this study were received by the investigators on 09/09/2025. Buffy coats were provided to the investigators in anonymized form, and the investigators had no access to donor-identifying information. All procedures involving human biological material were conducted in accordance with the principles of the Declaration of Helsinki.

## Data Availability

Bulk RNA-Seq data generated from sorted CRISPR TAMs, namely raw sequencing files and processed count matrices, are available at Gene Expression Omnibus (GEO) under accession number GSE346389. Additional data generated in this study are available from the corresponding author upon reasonable request.

## Competing Interests

The authors have declared that no competing interests exist.

## Funding

This work was funded by Fundação para a Ciência e a Tecnologia/Ministério da Educação, Ciência e Inovação (FCT/MECI, Portugal) through national funds to the Associate Laboratory LS4FUTURE (LA/P/0087/2020, doi.org/10.54499/LA/P/0087/2020), the iNOVA4Health Research Unit through the BRIDGE Project (UID/04462/2025, doi.org/10.54499/UID/04462/2025; UID/PRR/04462/2025, doi.org/10.54499/UID/PRR/04462/2025; UID/PRR2/04462/2025, doi.org/10.54499/UID/PRR2/04462/2025), LISBOA2030-FEDER-03115500, the RCD Project GlycoTAM (PTDC/BTM-TEC/0432/2021, doi.org/10.54499/PTDC/BTM-TEC/0432/2021), and the individual fellowship to GT (10.54499/2022/11642/BD). The authors also acknowledge funding from iBET (iBETxPlore Advanced Grant: PI-SinTra-767).

## Authors’ Contributions

Conceptualization: GT, GD, CB. Methodology: GT, GD, JE, CB. Formal analysis: GT, MP. Investigation: GT, MP.

Resources: CB.

Visualization: GT, MP.

Funding acquisition: CB. Supervision: GD, CB.

Writing – original draft: GT, MP, CB.

Writing – review C editing: GT, MP, GD, JE, CB.

## Supporting Information Captions

Supplementary Table S1. – Sequences of sgRNAs used for CRISPR-Cas9 gene editing.

Target gene, sgRNA identifier, targeted strand, protospacer adjacent motif (PAM), and sgRNA sequence are shown. The non-targeting control (NTC) and CD14 technical positive-control sgRNAs are also included.

Supplementary Table S2. – Primers used for PCR amplification and Sanger sequencing of CRISPR-targeted loci.

Forward and reverse primer sequences used to amplify regions surrounding each sgRNA target site are shown together with the primer used for Sanger sequencing.

Supplementary Table S3. – Antibodies used for flow-cytometry analysis.

Antibody target and fluorophore, supplier, catalogue number, and final staining concentration are indicated.

Supplementary Figure S1. Analytical and cellular assessment of nucleofection programs for CRISPR–Cas9 RNP editing in primary human monocytes. Primary human monocytes were nucleofected with pre-complexed CD14-targeting or non-targeting control (NTC) Cas9–sgRNA ribonucleoproteins using the CM-137, EA-100, or DK-100 nucleofection programs. A. Comparison of Sanger chromatogram deconvolution algorithms—TIDE (Tracking the Indels by Decomposition), ICE (Inference of CRISPR Edits), DECODR (Deconvolution of Complex DNA Repair), and SeqScreener—according to model fit (R²), estimated indel frequency, and predicted frameshift frequency for each nucleofection program. B. Representative phase-contrast microscopy images of monocytes nucleofected with NTC or CD14-targeting RNPs using CM-137, EA-100, or DK-100. Scale bar = 250 µm.

Supplementary Figure S2. Additional qualification of the primary-monocyte CRISPR–Cas9 RNP workflow. A. Representative flow-cytometry histograms of CD14 surface expression at day 7 following CD14-targeting or non-targeting control (NTC) RNP nucleofection in the presence or absence of the Alt-R Cas9 Electroporation Enhancer. Non-electroporated and electroporated controls are shown for comparison. B. Representative phase-contrast microscopy images of non-electroporated, electroporated-control, NTC, and CD14-targeted cultures in the presence or absence of electroporation enhancer. Scale bar = 250 µm. C. Representative flow-cytometry profiles of CD14, CD80, and CD206 expression in non-electroporated and CD14-targeted monocyte-derived macrophages following M0, M1/pro-inflammatory (IFN-γ + LPS), or M2/alternative-polarizing (IL-4 + IL-13) conditions. Data are from an exploratory single-donor experiment and are presented qualitatively.

Supplementary Figure S3. Screening and selection of sgRNAs targeting candidate lectins in HeLa S3 cells.

HeLa S3 cells were nucleofected with Cas9–sgRNA RNPs targeting CLEC4E, CLEC6A, CD209, SELL, or SIGLEC9, with the previously validated CD14 sgRNA included as a technical reference control. Genomic editing was assessed three days after nucleofection by Sanger sequencing and ICE analysis. Model fit (R²) and estimated indel frequencies are shown for the individual sgRNAs tested. Hatched bars indicate the sgRNA selected for each target for subsequent evaluation in primary human monocytes. Data represent a single technical screening experiment.

Supplementary Figure S4. Quality control of CRISPR–Cas9 editing, macrophage sorting and RNA integrity.

A. Total indel frequencies determined by ICE analysis in parallel day-3 monoculture samples following CRISPR–Cas9 RNP targeting of CLEC4E, CLEC6A, CD209, SELL, or SIGLEC9. B. ICE model-fit (R²) values for the corresponding Sanger chromatogram deconvolution analyses. For A–B, data are presented as mean ± SD from four independent monocyte donors, with individual donors shown. C. Representative post-sort analysis showing sequential gating of cells, single cells, viable cells, and CD45⁺ macrophages; the final CD45⁺ population showed 99.8% purity. D. RNA integrity number equivalent (RINe) of RNA isolated from sorted CD45⁺ macrophages across non-electroporated (NoEP), non-targeting control (NTC), and lectin-targeting conditions. Data are presented as mean ± SD from four independent monocyte donors, with individual donors shown.

Supplementary Figure S5. Complementary transcriptomic analyses following lectin perturbation in tumor-conditioned macrophages.

A–E. Pairwise principal component analyses (PCA) comparing non-targeting control (NTC) macrophages with A, CLEC4E; B, CLEC6A; C, CD209; D, SELL; and E, SIGLEC9-targeted macrophages. For visualization of condition-associated variation, the donor-associated component was removed while preserving experimental condition, and PCA was performed using the 500 most variable genes within each pairwise comparison. Points represent individual donor samples and shaded regions indicate 95% normal-data ellipses for each condition. F. Volcano plot comparing NTC and non-electroporated (NoEP) macrophages. Dotted lines indicate |log₂ fold change| = 1 and *p-value* = 0.001; genes meeting both exploratory thresholds are highlighted and labelled. G. PROGENy-inferred pathway-activity changes for NTC versus NoEP and each lectin-targeting condition versus NTC. Positive values indicate higher inferred pathway activity in the numerator condition and negative values indicate higher activity in the denominator condition. Dots indicate *p-value* < 0.05.

## Notes

### Competing Interest Statement

The authors have declared no competing interest.

