## Supplementary Tables for "CRISPR–Cas9 Editing of Myeloid Lectins Reveals Target-Dependent Transcriptional Remodeling in Tumor-Associated Macrophages"

Supplementary Table S1 – Sequences of sgRNAs used for CRISPR-Cas9 gene editing.

| Gene | | Strand | PAM | Sequence (5’ – 3’) |
|---|---|---|---|---|
| NTC | | – | – | UAACCGCGUAUACCGUUUAGA |
| CD14 | | - | TGG | GGGCTCAATGAACTCCGGCG |
| CLEC4E | #1 | - | AGG | GGTGGTTATCAACTCACAGG |
| | #3 | - | AGG | ACTGTCAGACCAGGTTGTCG |
| | #8 | + | GGG | AGGCACTGAGAAATAGGATG |
| | #10 | - | TGG | GAGCTCTCCTGCTACAATTA |
| | #15 | - | TGG | CGAGGGTCAGTGGCAATGGG |
| | #36 | - | TGG | AGAATTGTTGTCCATTGAAC |
| CLEC6A | #1 | + | GGG | GAGCAGAACTGTGTTGAGAT |
| | #2 | - | AGG | TCACAATGAAGCAAGCACTG |
| | #3 | + | AGG | CATATGGTGAAACTGGCAAA |
| | #4 | - | AGG | AGAAGCTGGGCAACATCCCC |
| | #11 | - | TGG | AGACCAAACCTTCTCTTCAC |
| | #12 | - | AGG | CAGAATGATTGGGCTCACCT |
| CD209 | #1 | - | AGG | CTTGGATTCCGACAGACTCG |
| | #2 | - | GGG | TCACCGCCTGCAAAGAAGTG |
| | #3 | - | AGG | CCATAAGTCAGGAACAATCC |
| | #12 | - | CGG | AGATCTACCAGGAGCTGACT |
| SELL | #1 | - | AGG | GGAAATGTCAGAGCACCCAG |
| | #2 | - | GGG | GATAGGAGGAATATGGACGT |
| | #12 | - | TGG | TTACACCTGCAACTGTGATG |
| | #17 | - | TGG | CACATCATGGAACCGACTGC |
| SIGLEC9 | #1 | + | TGG | TCGCATGGCTGGATTTACCC |
| | #2 | - | GGG | GGTAAATCCAGCCATGCGAG |
| | #22 | - | TGG | TGAGCCATTTCCCAAGACTG |
| | #30 | - | GGG | TCATGGTCAAGTTCTGAGGC |

Supplementary Table S2 – Primers used for PCR amplification and Sanger sequencing of CRISPR-targeted loci.

| Gene | | Forward Primer (5' – 3') | Reverse Primer (5' – 3') | Sanger Primer |
|---|---|---|---|---|
| CD14 | | AAGCCAGGCCTCAAGGTACT | GTTATCCACCTCGGGCAGCT | Rv |
| CLEC4E | #1 | GTGATTCCAAGGCTTCTGAC | TCCAACCCTCTTTTTCCCCT | Rv |
| | #3 | GAGGATCTCCTTTTCTGTTGAAGG | AGGAGAGAAGTGGGTGCTTG | Rv |
| | #8 | GTTTCCACCCTTTCTGCAATACA | AGTTCCCTGCTTGTTGAGGT | Rv |
| | #10 | GGAGAGTCTGCCTTAGAGCTTG | CCAGCTATATCTCACCTGTCCC | Fw |
| | #15 | GAGGATCTCCTTTTCTGTTGAAGG | AGGAGAGAAGTGGGTGCTTG | Rv |
| | #36 | GTGATTCCAAGGCTTCTGAC | TCCAACCCTCTTTTTCCCCT | Rv |
| CLEC6A | #1 | GCACTCACTCTTTTTTTTCA | GCTTTCAGGGATGGGGAAAT | Rv |
| | #2 | CTTCCTGGGGATTTTGGCTC | TGAGGCCATGATTCAGAAAA | Rv |
| | #3 | CTGTCCAAGAGCATGGAGCC | CTGGCACCTTTGTCCCTTCA | Rv |
| | #4 | ACTGGGTTTCTAGGTTCGTCT | TGAACACAACCAAATGTGCTCC | Fw |
| | #11 | GCACTCACTCTTTTTTTTCA | GCTTTCAGGGATGGGGAAAT | Rv |
| | #12 | ATAACCCCGCCAGCATAAAGG | AACATCATTCCAGCCCCATCC | Fw |
| CD209 | #1 | GGTGACATGAGTGACTCCAA | AAACTCAACCTCCTCAGGTC | Fw |
| | #2 | ACTCCTGGGTTCTCATGGGG | TCTGCAAAGTCATCTCTGAG | Fw |
| | #3 | CTTCTGTGCTGCCTTCTCCA | AAGCCCTGGTTCCAGATACTCAC | Fw |
| | #12 | CTTCTGTGCTGCCTTCTCCA | AAGCCCTGGTTCCAGATACTCAC | Rv |
| SELL | #1 | AATTCAGGGCAGTACCGATT | ATTGAGGGATTTTGTGGGGT | Fw |
| | #2 | CTCTAAGTGTCTAGCTCAGCC | ACCAGAAGCTGTTCTCATCC | Fw |
| | #12 | TAAAGCTTCTTGCCAGCCCT | AGAGCGGGAATCTACCATGC | Rv |
| | #17 | CTCTAAGTGTCTAGCTCAGCC | ACCAGAAGCTGTTCTCATCC | Fw |
| SIGLEC9 | #1 | CTCTGAGGAGGTCACTGTTC | TGTGCCTTACCTGTCACATT | Fw |
| | #2 | CTCTGAGGAGGTCACTGTTC | TGTGCCTTACCTGTCACATT | Fw |
| | #22 | ACTGTTCTCGATGAAGCGGG | CTCTGCAGGGAGACGTTCAG | Fw |
| | #30 | CCACGAACAAGACCGTCCAT | ACCCTCTTCTTGTTTGAGGCT | Fw |

Supplementary Table S3 – Antibodies used for flow-cytometry analysis.

| Antibody | Brand | Cat # | [Final] (μg/mL) |
|---|---|---|---|
| CD45 – BV421 | BD Biosciences | 563879 | 2.5 |
| CD45 – BV605 | BioLegend | 304042 | 2.5 |
| CD14 – BV711 | BioLegend | 301838 | 7.5 |
| CD64 – PE/Dazzle594 | BioLegend | 305032 | 5 |
| CD80 – APC | BioLegend | 305220 | 5 |
| CD86 – KiraviaBlue520 | BioLegend | 305452 | 5 |
| HLA-DR – BV421 | BioLegend | 307636 | 2.5 |
| CD163 – APC/Fire750 | BioLegend | 333634 | 10 |
| CD204 – PE/Cy7 | BioLegend | 371908 | 5 |
| CD206 – AF700 | BioLegend | 321132 | 7.5 |
| CLEC4E – PE/Cy5.5 | Novus Biologicals | NBP1-49311PECY55 | 10.6 |
| CLEC6A – PE/Cy5.5 | Novus Biologicals | FAB31141PECY55 | 8.3 |
| CD209 – PE/Cy7 | BioLegend | 330114 | 2.5 |
| SELL – PE/Cy7 | BD Biosciences | 565535 | 10 |
| SIGLEC9 – AF488 | R&D Systems | FAB1139G | 0.3 |
