## Supplementary Figures for "CRISPR–Cas9 Editing of Myeloid Lectins Reveals Target-Dependent Transcriptional Remodeling in Tumor-Associated Macrophages"

### Supplementary Figure S1

## A

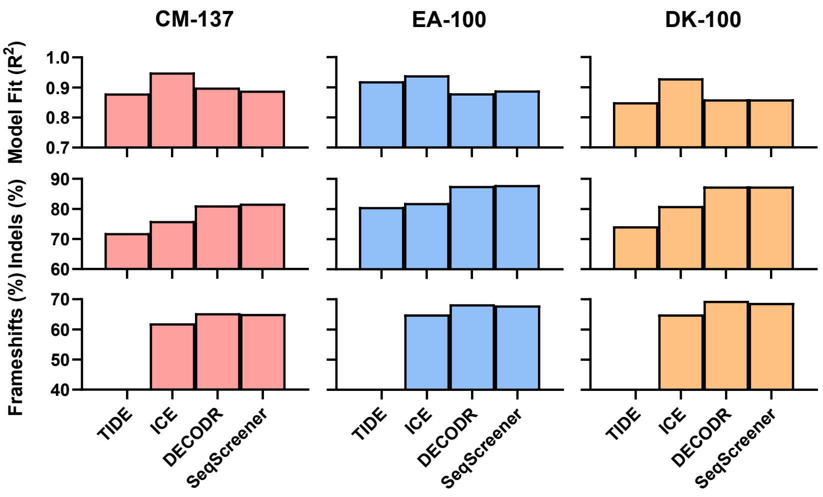


## B

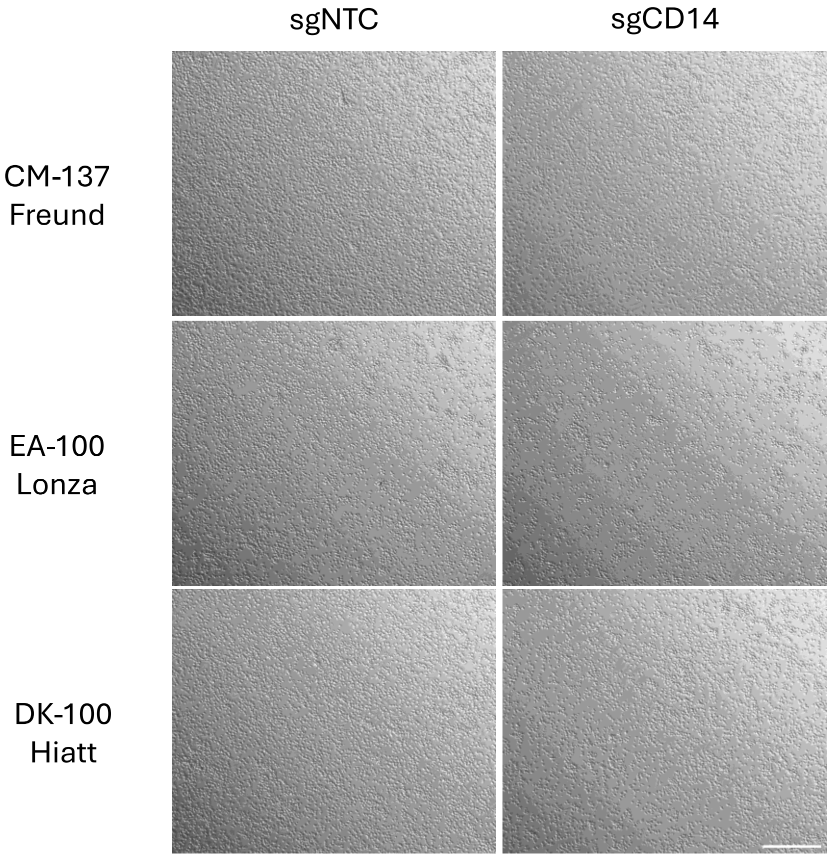

### Supplementary Figure S1

A

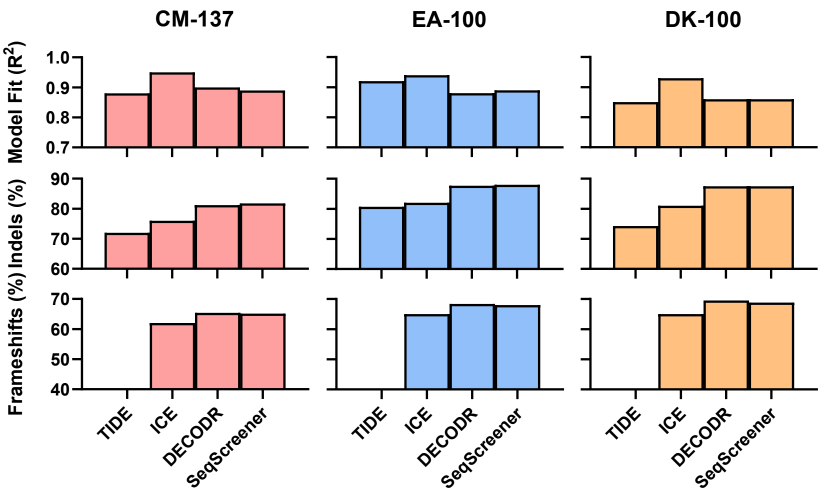


B

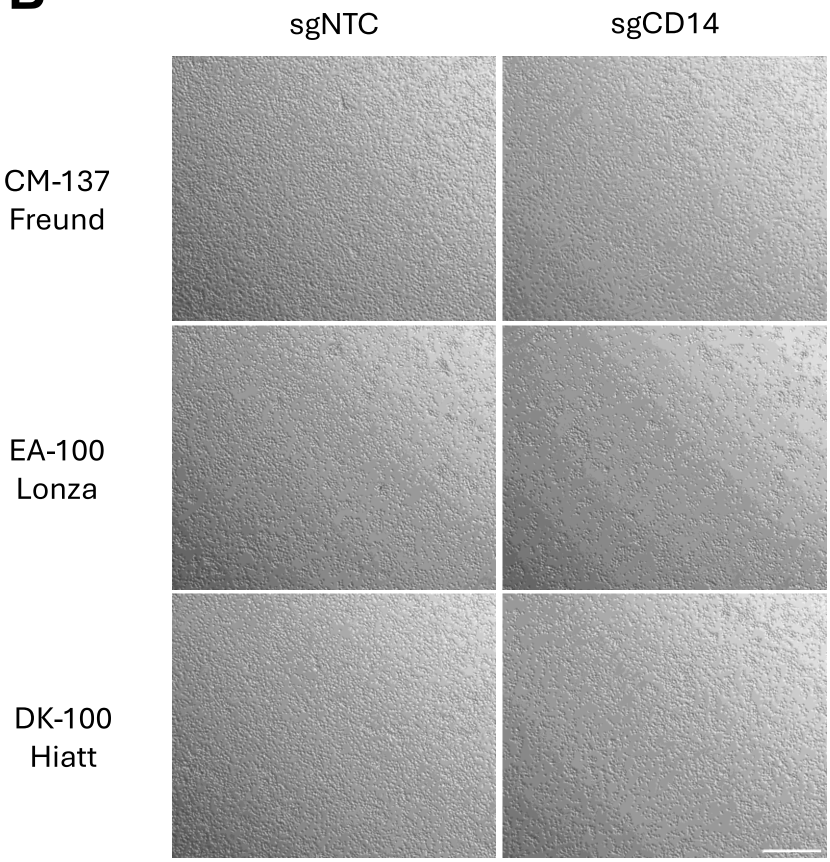

#### Supplementary Figure S2

**A**

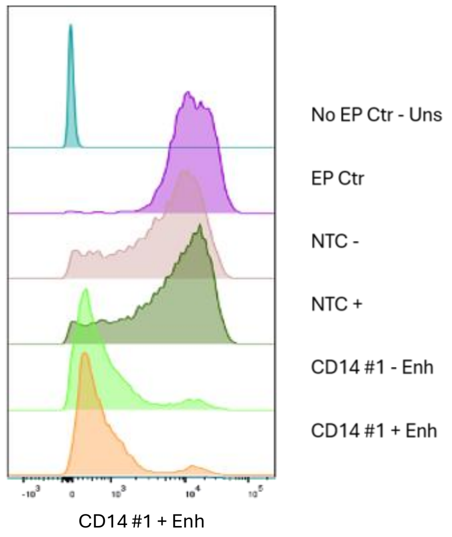


**B**

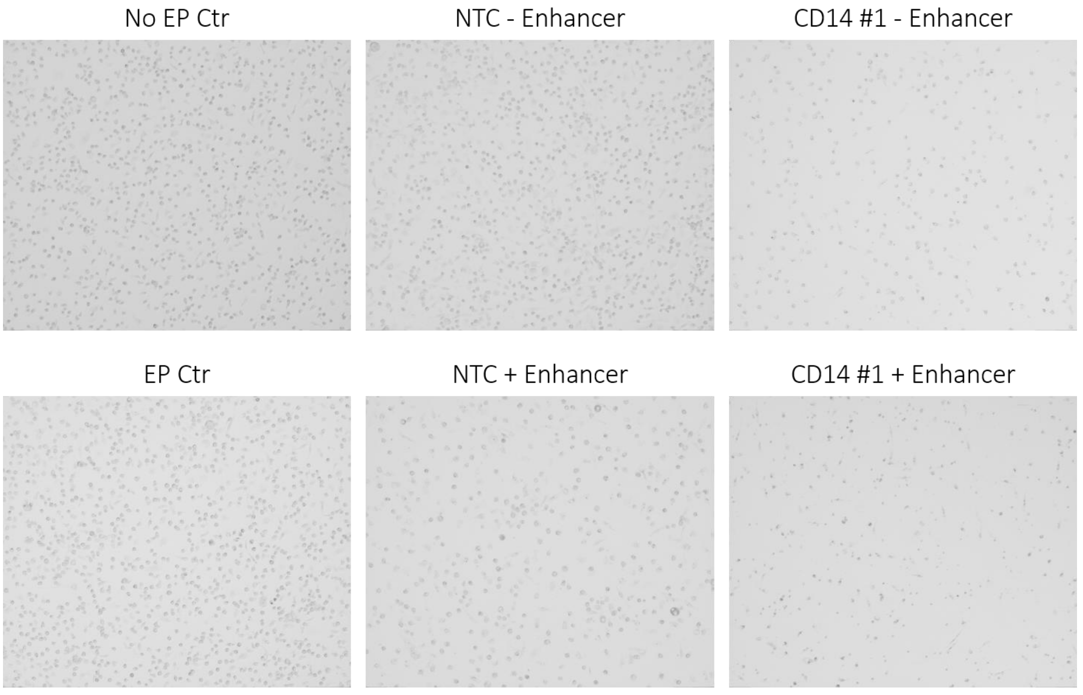


**C**

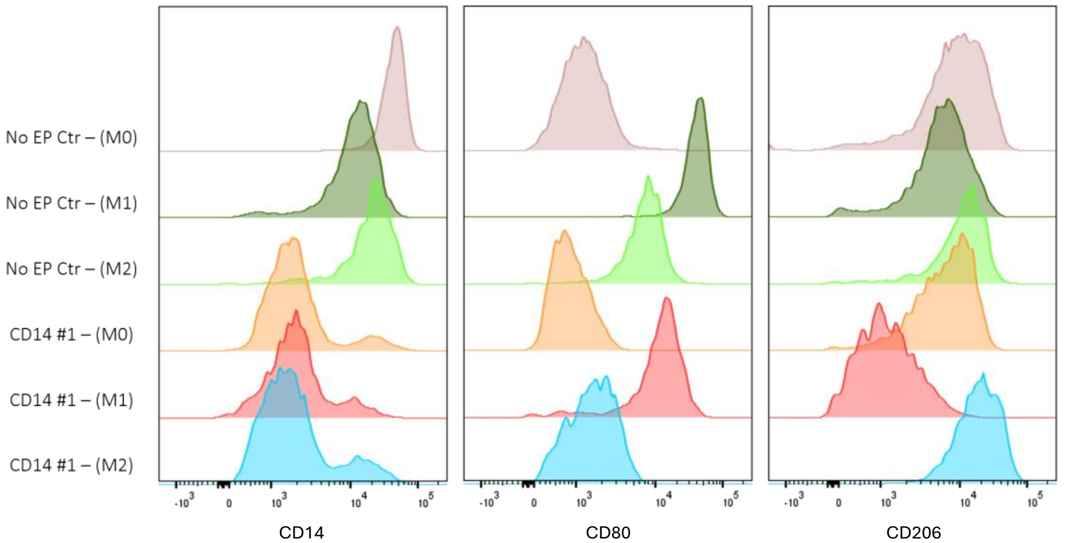

### Supplementary Figure S2

A

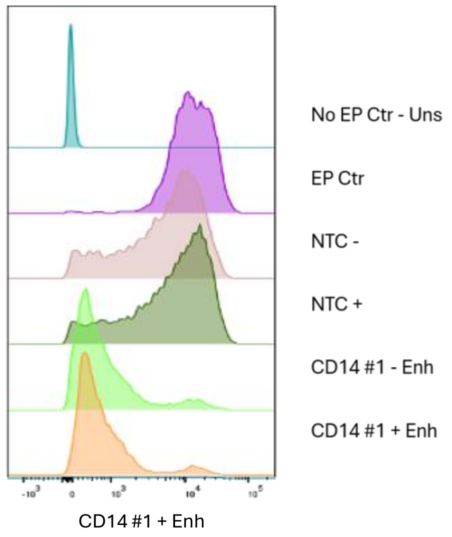


B

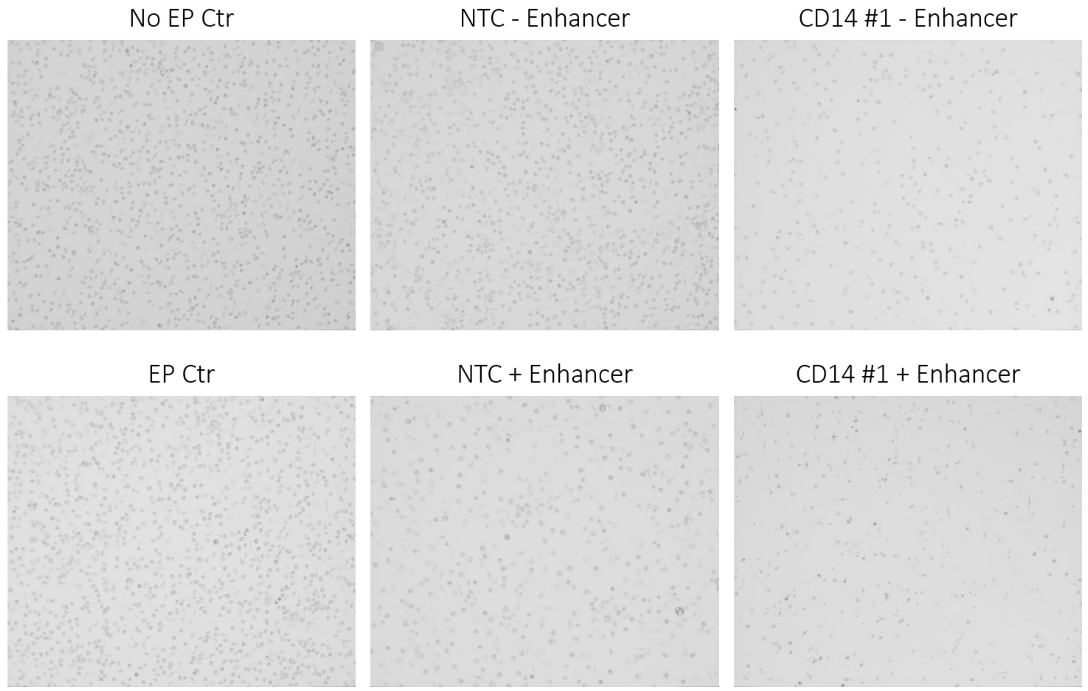


C

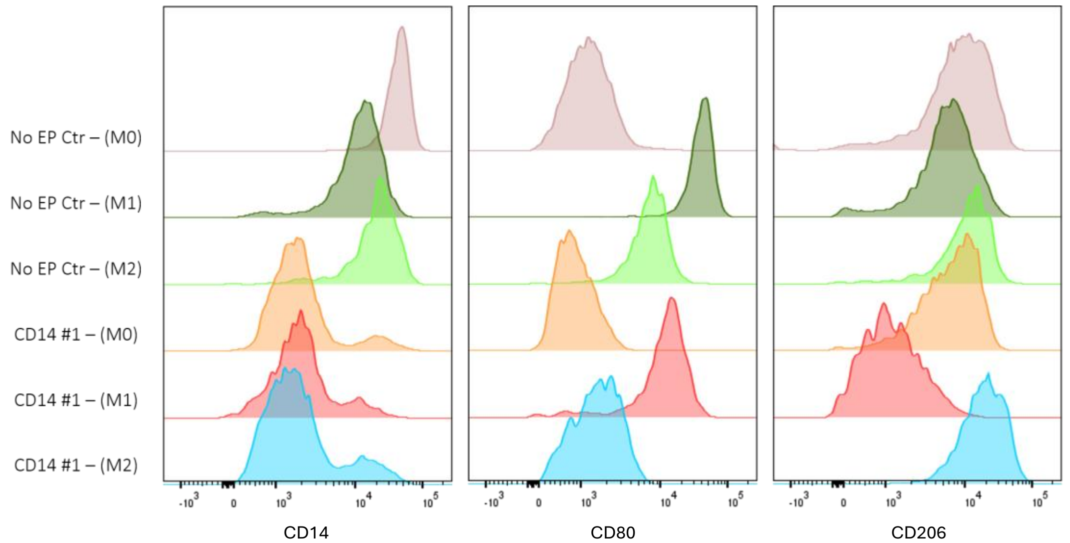

### Supplementary Figure S3

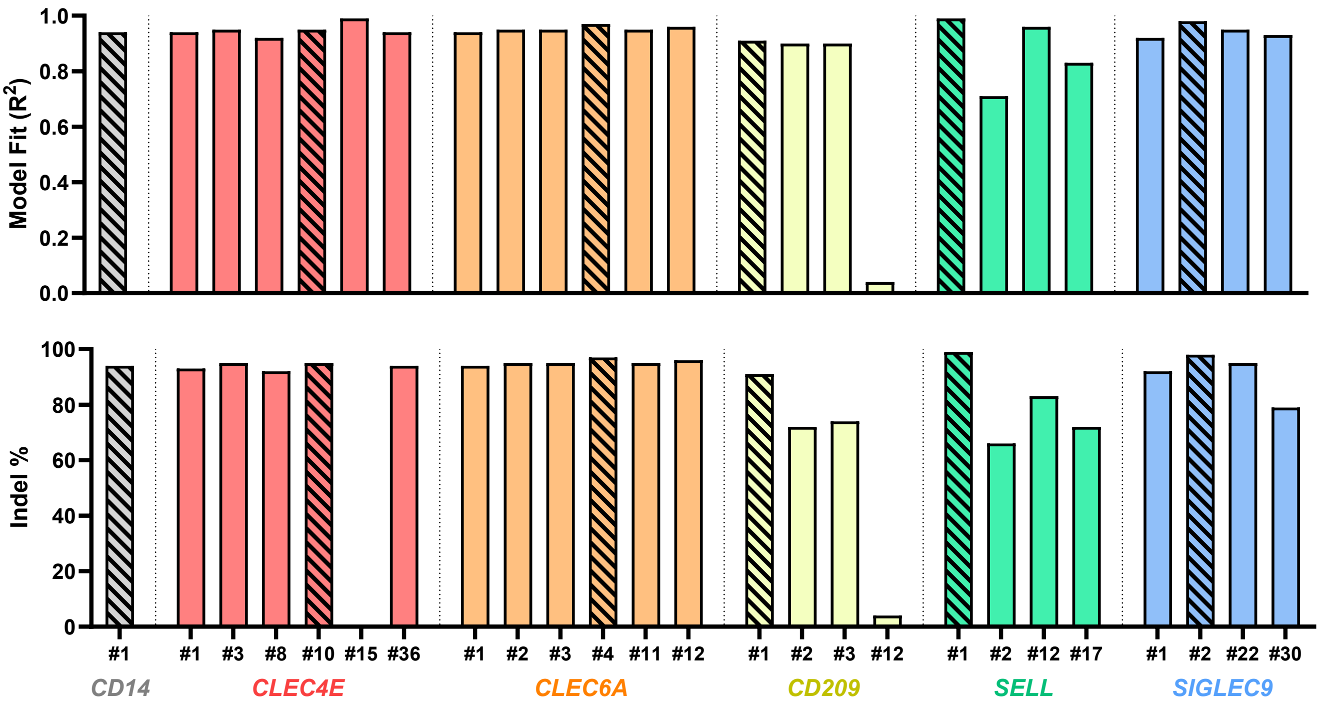

### Supplementary Figure S4

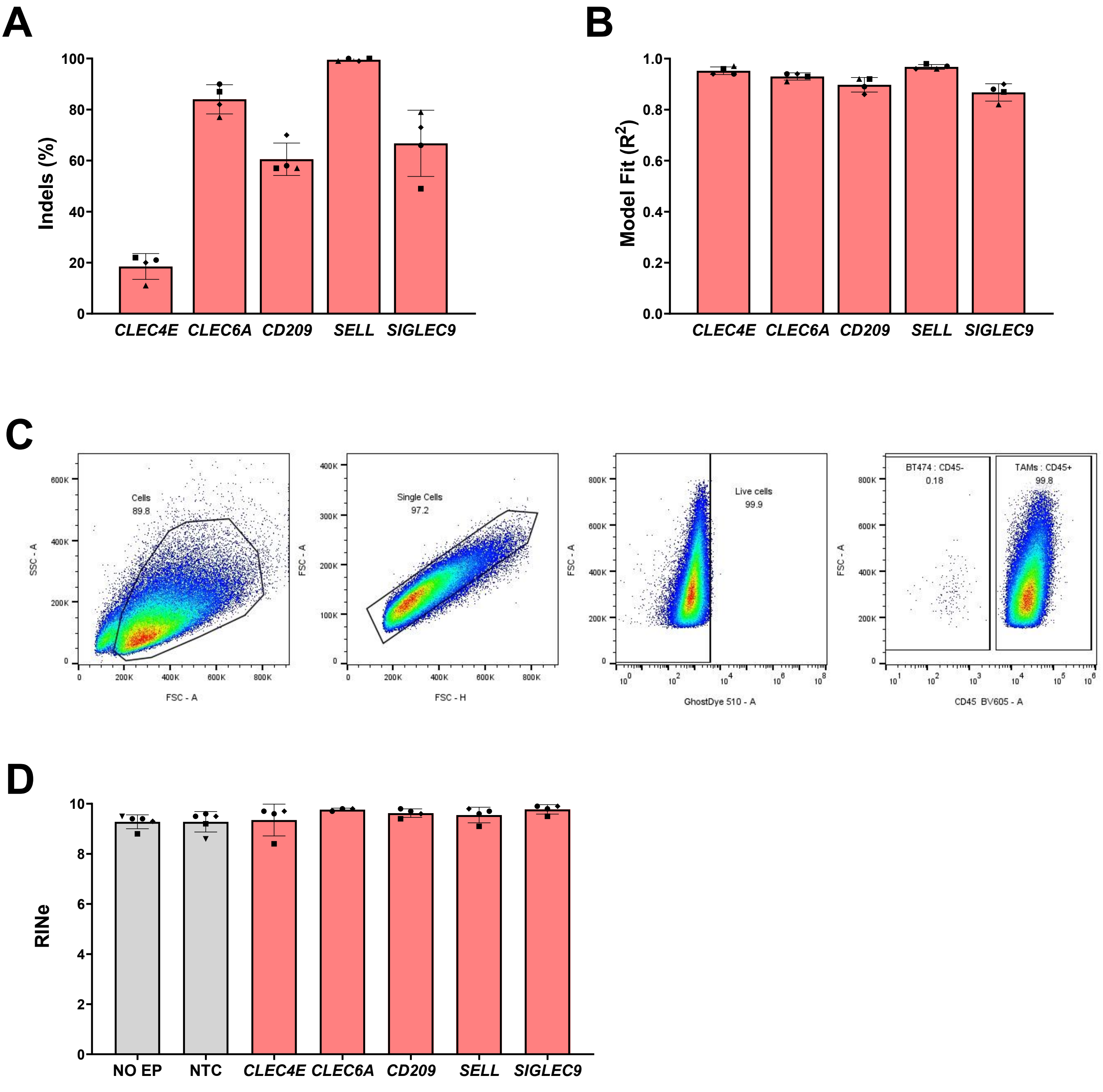

### Supplementary Figure S5

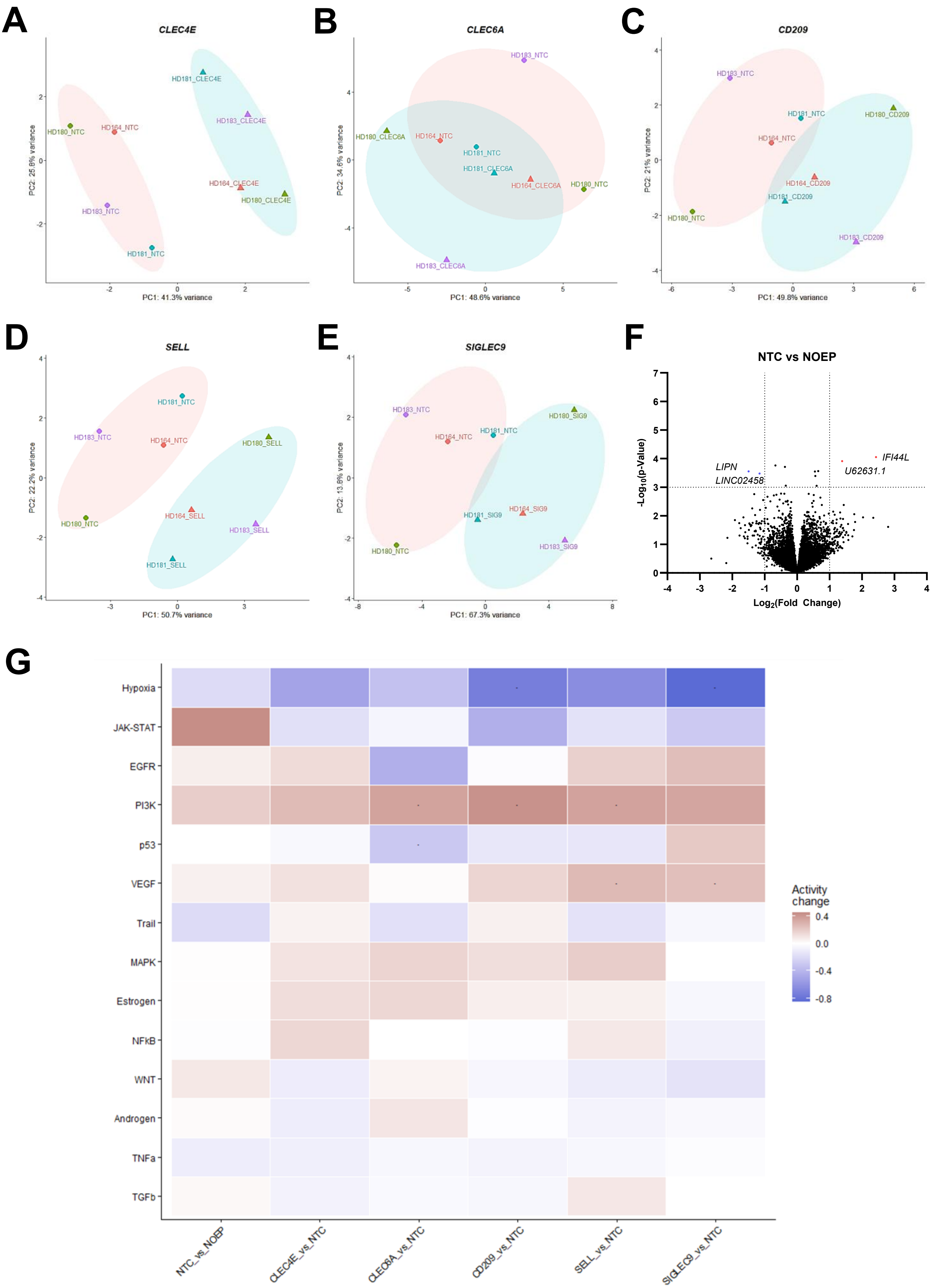
